# VADR: validation and annotation of virus sequence submissions to GenBank

**DOI:** 10.1101/852657

**Authors:** Alejandro A Schäffer, Eneida L Hatcher, Linda Yankie, Lara Shonkwiler, J Rodney Brister, Ilene Karsch-Mizrachi, Eric P Nawrocki

## Abstract

**Background:** GenBank contains over 3 million viral sequences. The National Center for Biotechnology Information (NCBI) previously made available a tool for validating and annotating influenza virus sequences that is used to check submissions to GenBank. Before this project, there was no analogous tool in use for non-influenza viral sequence submissions.

**Results:** We developed a system called VADR (Viral Annotation DefineR) that validates and annotates viral sequences in GenBank submissions. The annotation system is based on the analysis of the input nucleotide sequence using models built from curated RefSeqs. Hidden Markov models are used to classify sequences by determining the RefSeq they are most similar to, and feature annotation from the RefSeq is mapped based on a nucleotide alignment of the full sequence to a covariance model. Predicted proteins encoded by the sequence are validated with nucleotide-to-protein alignments using BLAST. The system identifies 43 types of “alerts” that (unlike the previous BLAST-based system) provide deterministic and rigorous feedback to researchers who submit sequences with unexpected characteristics. VADR has been integrated into GenBank’s submission processing pipeline allowing for viral submissions passing all tests to be accepted and annotated automatically, without the need for any human (GenBank indexer) intervention. Unlike the previous submission-checking system, VADR is freely available (https://github.com/nawrockie/vadr) for local installation and use. VADR has been used for Norovirus submissions since May 2018 and for Dengue virus submissions since January 2019. Other viruses with high numbers of submissions will be added incrementally.

**Conclusion:** VADR improves the speed with which non-flu virus submissions to GenBank can be checked and improves the content and quality of the GenBank annotations. The availability and portability of the software allow researchers to run the GenBank checks prior to submitting their viral sequences, and thereby gain confidence that their submissions will be accepted immediately without the need to correspond with GenBank staff. Reciprocally, the adoption of VADR frees GenBank staff to spend more time on services other than checking routine viral sequence submissions.

## Background

As of September 2019, GenBank [1] contained more than 3 million viral sequences totaling over 4 billion nucleotides in length and including over 180,000 complete genomes for viruses other than influenza. More than 250,000 of these sequences were submitted in 2018. All sequence submissions are validated prior to deposition in GenBank. Automated validation and annotation methods become increasingly important as sequence submission numbers grow.

Table 1 shows the number of sequences for the 16 virus species with the most sequences in GenBank. Influenza sequences are the second most abundant and the National Center of Biotechnology Information (NCBI), where GenBank is housed, has expended considerable effort to organize flu sequences and streamline the submission of new influenza virus sequences, including a tool to validate and annotate flu submissions called FLAN [2]. The influenza virus sequence submission tool (https://www.ncbi.nlm.nih.gov/genome/viruses/variation/help/flu-help-center/submit-flu-sequences/) is implemented specifically for influenza with many hard-coded features. It has proven impractical to reuse the influenza virus submission code for other viruses.

**Table 1.** Viruses with the highest number of sequences in GenBank as of October 10, 2019. The number of sequences for the segmented viruses Influenza and Rotavirus are the sums over all their segments.

| species | #seqs | family |
| --- | --- | --- |
| HIV-1 | 850,115 | *Retroviridae* |
| Influenza A virus | 684,026 | *Orthomyxoviridae* |
| Hepacivirus C | 244,533 | *Flaviviridae* |
| Hepatitis B virus | 114,306 | *Hepadnaviridae* |
| Influenza B virus | 100,373 | *Orthomyxoviridae* |
| Rotavirus A | 73,375 | *Reoviridae* |
| SIV | 44,374 | *Retroviridae* |
| Norovirus (Norwalk virus) | 40,925 | *Caliciviridae* |
| Enterovirus A | 31,478 | *Picornaviridae* |
| PRRSV | 29,081 | *Arteriviridae* |
| Dengue virus | 28,564 | *Flaviviridae* |
| Human orthopneumovirus | 24,384 | *Pneumoviridae* |
| Enterovirus B | 23,865 | *Picornaviridae* |
| Rabies lyssavirus | 23,771 | *Rhabdoviridae* |
| West Nile virus | 21,563 | *Flaviviridae* |
| Measles morbillivirus | 17,233 | *Paramyxoviridae* |

NCBI’s Virus Variation Resource [3] (https://www.ncbi.nlm.nih.gov/genome/viruses/variation) includes specialized components that attempt to normalize annotation of previously submitted sequences for rotaviruses, dengue virus, West Nile virus, ebolaviruses, Zika virus, and MERS coronavirus. The resulting standardized annotation supports virus-specific searches using gene and protein names. However, the Virus Variation Resource does not support the submission of new sequences, and its tools are not generalizable, such that creating components for additional virus species is laborious.

### Automating validation of sequence submissions

Early in the history of GenBank, Michael Waterman presciently wrote that “Entering new sequences into the databases requires the database staff to analyze and interpret the sequences and the associated scientific literature [4].” Even earlier, Margaret Dayhoff had justified her enormous efforts at sequence database curation reasoning that “a carefully verified collection [of sequences] was more economical in the long run than a quick and dirty collection” [5]. At NCBI, sequence submissions have been checked by indexers since NCBI began operating GenBank, but over the years, limitations to this arrangment have become apparent. First, the number and size of sequence submissions has grown much faster than the NCBI budget and the capability to hire more indexers. Second, although GenBank indexers are trained rigorously and homogeneously and use the same software, there is no formal mechanism to enforce “inter-observer agreement”, meaning that all GenBank indexers would be guaranteed to give the same evaluation of the same submission. Third, researchers who wish to submit their sequences cannot reproduce all the checks that GenBank indexers do. Consequently, problematic sequences cause delays and e-mail interactions between submitters and indexers that might be avoidable using a more deterministic and open system of checks. The importance of transparency in curatorial analysis was emphasized by Walter Goad, one of the founders of GenBank: “It is important that we be perceived by the molecular biology community as offering free and open access to the information *and programs* we will be collecting” [5].

For bacterial genomes, these limitations have been addressed by the release of NCBI’s Prokaryotic Genome Annotation Pipeline (PGAP) [6], which is now available for download as a standalone package (https://github.com/ncbi/pgap/releases). For influenza virus, the limitations were addressed first by FLAN [2] and then by the influenza virus submission tools provided by GenBank [3]. Tools for checking submissions of non-influenza viruses have lagged but in the work described herein we describe new software that begins to close the gap. Importantly, our new software can already be used for many viruses (currently excluding circular genome and segmented viruses) breaking from the paradigm of developing code for one virus at a time that has limited the breadth of the Virus Variation Resource.

### Software for viral genome annotation

Several software packages exist for viral genome annotation. VAPiD [7] is designed to simplify the submission of complete viral genome sequences to GenBank by adding metadata, and annotating each input sequence based on comparison with its best-matching reference sequence in a large reference dataset derived from GenBank. VIGOR [8, 9] annotates input sequences by first identifying the most relevant reference database in its collection and then comparing all reference protein and mature peptide sequences in that database to the input sequence to determine its annotation and to identify certain types of errors. Both programs detect and report some types of unexpected errors, such as premature stop codons.

Additional programs for viral annotation include VGAS [10], which incorporates *ab initio* ORF prediction as well as similarity-based annotation, and VIGA [11] which can be optimized for speed for huge metagenomics datasets. All of these programs focus primarily on annotating protein-coding regions, although VIGA additionally identifies some types of RNAs and CRISPR repeat elements. Among these four programs, VAPiD and VIGOR provide some capabilities to prepare submissions for GenBank and hence are closer to solving the submission checking problem considered here than are VGAS and VIGA.

### VADR: Viral Annotation DefineR

In this work, we describe the design and implementation of a new reference-based general software tool called VADR (Viral Annotation DefineR), for the validation and annotation of virus sequences from characterized species that have an open (non-circular) genome less than 25Kb in length and for which a reference (e.g. RefSeq entry) exists. We discuss the use of VADR in evaluating GenBank submissions of norovirus and dengue virus sequences. We compare VADR performance with that of VAPiD and VIGOR and show how it identifies more potential problems with sequences, making it more appropriate for use by GenBank indexers interested in manually reviewing those problems prior to sequence deposition.

VADR compares each input sequence to a library of homology models of viral species built from reference sequences from the RefSeq database [12], identifies the most similar model, and uses that model to compute an alignment to the RefSeq from which feature annotation boundaries (e.g. coding sequences (denoted CDS), mature peptides, ncRNAs) are derived. Finally, CDS features that encode proteins are validated for protein-coding potential using *blastx*. Submitted sequences that are confidently aligned and annotated with VADR *pass* and are cleared for automatic entry into GenBank. In contrast, when a submitted sequence is evaluated by VADR and the comparison to its matching RefSeq reveals the input sequence is divergent in various ways (e.g. early stop codon, regions of low nucleotide similarity), then the sequence *fails*. Failure means that the sequence is flagged for manual review by an NCBI expert curator, called a “GenBank indexer”, and the sequence is prevented from automatic entry into GenBank. If all sequences in a submission pass, all sequences will automatically be deposited into GenBank. If at least one sequence fails, a report with sequence-specific errors is generated and reported to the submitter or reviewed by the indexer, who can clear sequences for submission or contact the submitter for further investigation of the apparent problems.

VADR’s output annotation of each sequence is a five-column feature table conforming to the GenBank feature table syntax rules, and includes all desired features from the RefSeq that could be mapped onto the input sequence via sequence alignment. These features include protein coding regions, the “mature peptide” cleavage products of proteins, genes, noncoding RNAs, and structural RNA features. In the nomenclature of the International Nucleotide Sequence Database Collaboration (INSDC), these are denoted as *CDS*, *mat peptide*, *gene*, *ncRNA*, and *stem loop* features, respectively. Submitters rarely annotate RNA features in GenBank virus records even though many viruses have functionally important non-coding RNAs (ncRNAs) and other structural RNA features [13, 14]. VADR’s capability to annotate these features adds value to the resulting GenBank records.

VADR is in production usage for sequence submissions of norovirus and dengue virus, two prominent human pathogens from Table 1. Dengue virus is a member of family *Flaviviridae*, which also includes the West Nile Virus and Hepacivirus C (Hepatitis C). (We use upper case Norovirus and Dengue virus when referring to formal names and lower case norovirus and dengue virus when referring to attributes of sequences.) VADR is publicly available, so that researchers who plan to submit norovirus or dengue virus sequences to GenBank may pre-check their submissions with VADR to determine which sequences pass and fail.

## Implementation

VADR is written in Perl and is available at https://github.com/nawrockie/vadr. The new software uses the existing software packages and libraries listed in Table 2.

**Table 2.**
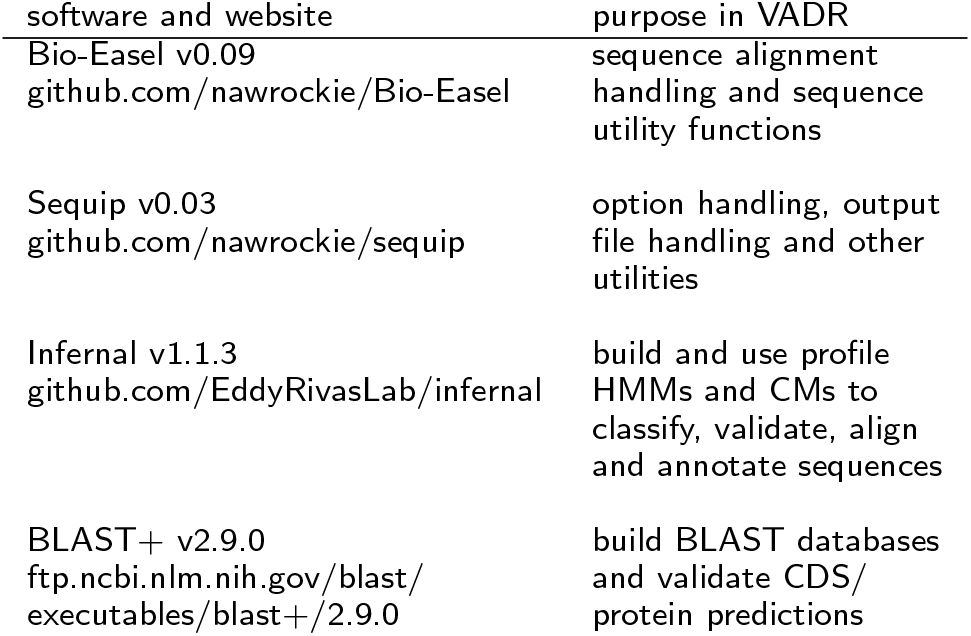
Software packages and libraries used within VADR.

VADR contains two major scripts. The *v-build.pl* script is used to build models of a viral species (e.g. norovirus). The *v-annotate.pl* script uses those models to annotate sequences from the viral species represented by the models. The main workflow of VADR is depicted in Figure 1.

**Figure 1.**
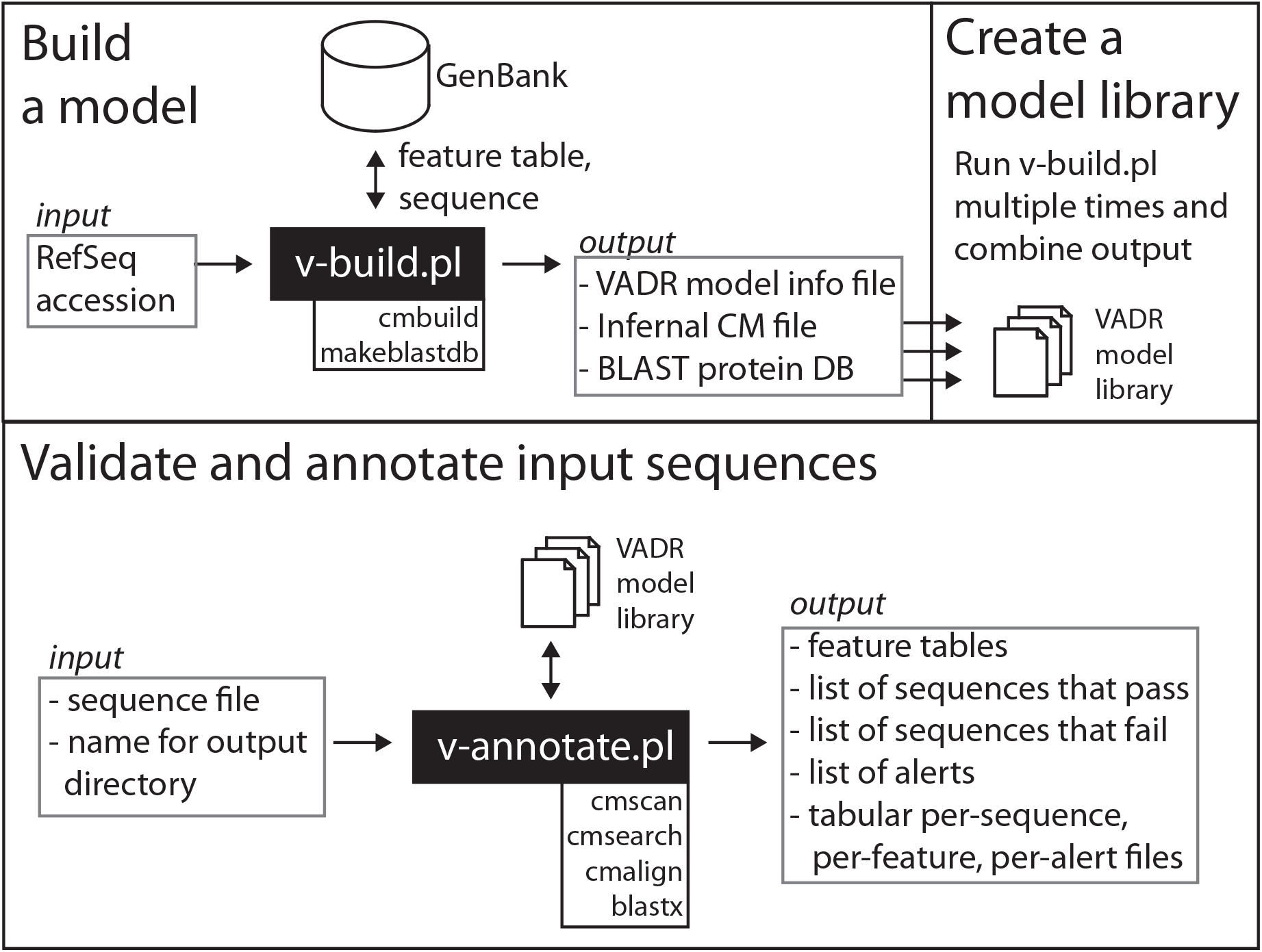
VADR workflow schematic illustrating uses of the two main VADR scripts. *v-build.pl* can be used once to build a single model or repeatedly to build a library of models. *v-annotate.pl* can be used with a model or model library to validate and annotate input sequences.

### The build module: *v-build.pl*

The *v-build.pl* script takes as input two arguments: the RefSeq accession to be modeled (e.g. “NC 001959”, for Norovirus genotype GI) and the name of the output directory to create and to populate with output files.

The script first retrieves a FASTA-formatted sequence file and five-column feature table files for the RefSeq and any protein sequences associated with CDS features of the RefSeq from NCBI. *v-build.pl* parses these files and outputs a VADR “model information” file with coordinates of CDS, mat peptide, and gene features and product and exception qualifiers. By default, all other feature types and qualifiers are ignored, although command-line options can be used to specify that additional feature types, such as ncRNA and stem loop, and qualifiers should be included. *v-build.pl* also outputs a covariance model (CM) file for the input RefSeq created using Infernal v1.1.3’s *cmbuild* program. If the RefSeq nucleotide sequence being modeled has known secondary structural elements, such as stem loops, then a Stockholm format file with structure annotation can be provided, and the resulting CM file will model the specified structure. This structure will inform the sequence-and-structure based alignment of input sequences by Infernal’s *cmalign* program in the annotation stage of *v-annotate.pl*. By default, if no Stockholm file is input to *v-build.pl*, then the RefSeq sequence is modeled without secondary structure. Additionally, *v-build.pl* uses the *makeblastdb* program from BLAST v2.9.0+ [15] to create a BLAST database from amino acid translations of the RefSeq CDS features. *v-annotate.pl* uses this database with *blastx* to validate its nucleotide-based predictions of CDS features. This design in which VADR annotates nucleotide sequences, but validates with protein alignments allows us to mitigate the known limitation that the 4-letter nucleic acid alphabet is less sensitive in homology searching than the 20-letter amino acid alphabet [16], while retaining the capability to annotate ncRNAs and other features that are not translated.

*v-build.pl* is designed to allow input alignments instead of single RefSeq accessions. Although models built from alignments have not yet been tested, we plan to use alignment-derived models in the future.

### VADR model library

VADR includes a library of models generated by *v-build.pl*, represented by a “vadr.minfo” model info file, a “vadr.cm” CM file, and BLAST database files that include information and models for 194 viral RefSeq genomes (v1.0 VADR library). These 194 RefSeqs comprise 9 norovirus sequences, 4 dengue virus sequences, and an additional 181 RefSeqs from other viruses in the families *Caliciviridae* and *Flaviviridae*. The additional 181 RefSeqs allow VADR to recognize when an input sequence purported to be norovirus or dengue virus is in fact a different *Caliciviridae* or *Flaviviridae* taxon. This library is currently used by GenBank indexers on submitted sequences to annotate norovirus and dengue virus sequences and to recognize some taxonomically misclassified sequences. Users are able to build their own model libraries with *v-build.pl* to use instead of this default library. We plan to add additional models to the default set in future versions after testing them internally at NCBI.

### The annotation module: *v-annotate.pl*

The *v-annotate.pl* script takes as input a sequence file in FASTA format and the name of an output directory. It uses the models in the default v1.0 VADR model library to analyze and to annotate the input sequences. *v-annotate.pl* creates output files including a feature table with annotations in the output directory. The workflow of the *v-annotate.pl* script consists of four stages: classification, coverage determination, alignment-based annotation and protein validation, each of which is described in more detail below.

During each stage, specific possible problems are identified and reported in the output as “alerts”. An alert is meant to inform a VADR user about an unusual, unexpected, or otherwise remarkable characteristic of a sequence. There are 43 types of alerts listed and briefly explained in Table 3. Alerts can pertain to either the entire sequence being evaluated (indicated by “S” in the “S/F” column in Table 3), or a specific feature of that sequence (indicated by “F” in the “S/F” column). By default, 38 of the 43 alert types are *fatal* in that they cause a sequence to fail annotation, whereas 5 alerts are reported but not fatal. A sequence passes if and only if it generates zero fatal alerts. Nearly all alert types can be set as fatal or not fatal using the command line options *--alt pass* and *--alt fail* to *v-annotate.pl*, but four alert types are always fatal and cannot be changed: *noannotn*, *revcompl*, *unexdivg*, and *noftrann*.

**Table 3.** Attributes of the 43 types of VADR alerts organized by the stage in which they are detected. The S/F column indicates whether the alert applies to an entire sequence (S) or to one feature (F) in a sequence. The five non-fatal alerts (four detected in the classification stage, and one in the coverage stage) do not cause a sequence to fail and are not reported in the output feature table. Codes marked with * are always fatal; all other codes can be set to fatal or non-fatal with command-line options to v-annotate.pl.

| code | S/F | error message | description |
| --- | --- | --- | --- |
| **Fatal alerts detected in the classification stage** |  |  |  |
| noannotn* | S | NO_ANNOTATION | no significant similarity detected |
| revcompl* | S | REVCOMPLEM | sequence appears to be reverse complemented |
| incsbgrp | S | INCORRECT_SPECIFIED_SUBGROUP | score difference too large between best overall model and best specified subgroup model |
| incgroup | S | INCORRECT_SPECIFIED_GROUP | score difference too large between best overall model and best specified group model |
| **Non-fatal alerts detected in the classification stage** |  |  |  |
| qstsbgrp | S | QUESTIONABLE_SPECIFIED_SUBGROUP | best overall model is not from specified subgroup |
| qstgroup | S | QUESTIONABLE_SPECIFIED_GROUP | best overall model is not from specified group |
| indfclas | S | INDEFINITE_CLASSIFICATION | low score difference between best overall model and second best model (not in best model's subgroup) |
| lowscore | S | LOW_SCORE | score to homology model below low threshold |
| **Fatal alerts detected in the coverage stage** |  |  |  |
| lowcovrg | S | LOW_COVERAGE | low sequence fraction with significant similarity to homology model |
| dupregin | S | DUPLICATE_REGIONS | similarity to a model region occurs more than once |
| discontn | S | DISCONTINUOUS_SIMILARITY | not all hits are in the same order in the sequence and the homology model |
| indfstrn | S | INDEFINITE_STRAND | significant similarity detected on both strands |
| lowsim5s | S | LOW_SIMILARITY_START | significant similarity not detected at 5' end of the sequence |
| lowsim3s | S | LOW_SIMILARITY_END | significant similarity not detected at 3' end of the sequence |
| lowsimis | S | LOW_SIMILARITY | internal region without significant similarity |
| **Non-fatal alerts detected in the coverage stage** |  |  |  |
| biasdseq | S | BIASED_SEQUENCE | high fraction of score attributed to biased sequence composition |
| **Fatal alerts detected in the annotation stage** |  |  |  |
| unexdivg* | S | UNEXPECTED_DIVERGENCE | sequence is too divergent to confidently assign nucleotide-based annotation |
| noftrann* | S | NO_FEATURES_ANNOTATED | sequence similarity to homology model does not overlap with any features |
| mutstart | F | MUTATION_AT_START | expected start codon could not be identified |
| mutendcd | F | MUTATION_AT_END | expected stop codon could not be identified, predicted CDS stop by homology is invalid |
| mutendns | F | MUTATION_AT_END | expected stop codon could not be identified, no in-frame stop codon exists 3' of predicted valid start codon |
| mutendex | F | MUTATION_AT_END | expected stop codon could not be identified, first in-frame stop codon exists 3' of predicted stop position |
| unexleng | F | UNEXPECTED_LENGTH | length of complete coding (CDS or mat_peptide) feature is not a multiple of 3 |
| cdsstopn | F | CDS_HAS_STOP_CODON | in-frame stop codon exists 5' of stop position predicted by homology to reference |
| peptrans | F | PEPTIDE_TRANSLATION_PROBLEM | mat_peptide may not be translated because its parent CDS has a problem |
| pepadjcy | F | PEPTIDE_ADJACENCY_PROBLEM | predictions of two mat_peptides expected to be adjacent are not adjacent |
| indfantn | F | INDEFINITE_ANNOTATION | nucleotide-based search identifies CDS not identified in protein-based search |
| indf5gap | F | INDEFINITE_ANNOTATION_START | alignment to homology model is a gap at 5' boundary |
| indf5loc | F | INDEFINITE_ANNOTATION_START | alignment to homology model has low confidence at 5' boundary |
| indf3gap | F | INDEFINITE_ANNOTATION_END | alignment to homology model is a gap at 3' boundary |
| indf3loc | F | INDEFINITE_ANNOTATION_END | alignment to homology model has low confidence at 3' boundary |
| lowsim5f | F | LOW_FEATURE_SIMILARITY_START | region within annotated feature at 5' end of sequence lacks significant similarity |
| lowsim3f | F | LOW_FEATURE_SIMILARITY_END | region within annotated feature at 3' end of sequence lacks significant similarity |
| lowsimif | F | LOW_FEATURE_SIMILARITY | region within annotated feature lacks significant similarity |
| **Fatal alerts detected in the protein validation stage** |  |  |  |
| cdsstopp | F | CDS_HAS_STOP_CODON | stop codon in protein-based alignment |
| indfantp | F | INDEFINITE_ANNOTATION | protein-based search identifies CDS not identified in nucleotide-based search |
| indf5plg | F | INDEFINITE_ANNOTATION_START | protein-based alignment extends past nucleotide-based alignment at 5' end |
| indf5pst | F | INDEFINITE_ANNOTATION_START | protein-based alignment does not extend close enough to nucleotide-based alignment 5' endpoint |
| indf3plg | F | INDEFINITE_ANNOTATION_END | protein-based alignment extends past nucleotide-based alignment at 3' end |
| indf3pst | F | INDEFINITE_ANNOTATION_END | protein-based alignment does not extend close enough to nucleotide-based alignment 3' endpoint |
| indfstrp | F | INDEFINITE_STRAND | strand mismatch between protein-based and nucleotide-based predictions |
| insertnp | F | INSERTION_OF_NT | too large of an insertion in protein-based alignment |
| deletinp | F | DELETION_OF_NT | too large of a deletion in protein-based alignment |

One motivation for reporting alerts is to identify sequences that should be reviewed by a GenBank indexer as opposed to being entered automatically into the GenBank database without any human inspection. Sequences that pass are automatically entered into GenBank, while sequences that fail are not and must be manually reviewed by an indexer. The alert output contains specific information (e.g. nucleotide positions) when possible to facilitate further inspection by the indexer or by the submitter running VADR locally.

#### Classification stage

Each *v-annotate.pl* input query sequence is first scored against each model in the library, which by default is the VADR v1.0 library of 194 models. The comparison is made between HMMs from the CM file and each sequence using an abbreviated version of the HMMER3 [17, 18] filter pipeline as implemented in Infernal v1.1.3 [19]. The full HMMER3 pipeline includes more than five stages, but only the first three stages are carried out during this classification stage of *v-annotate.pl*, which computes a local forward score of a window of the input sequence. The window boundaries reported at this stage of the filter pipeline are inexact but this imprecision is irrelevant for the classification stage’s goal of identifying the model that scores highest for each input sequence. The window boundaries are more precisely defined at later stages. In practice, the window reported at this stage will often be the entire sequence. For each sequence *S*, the highest scoring window and associated model *M* are found, and *M* (*S*) is defined as the best-scoring model for *S* to be used in the subsequent stages.

#### Coverage determination stage

For each sequence *S*, the model *M* (*S*) is used to determine the sequence coverage. This is achieved again using the HMMER3 filter pipeline as implemented in Infernal 1.1.3, but now using the full pipeline that reports local matches in each input sequence, with more precise endpoints than the abbreviated pipeline used in the classification stage. For sequences that include internal short stretches of sequence that are dissimilar from the RefSeq, multiple matches may be returned at this stage, with the dissimilar regions not covered by any of the matches. The coverage of *S* is determined as the fraction of nucleotides in *S* that occur in any of the alignments on the top strand (+ strand). An alert (*lowcovrg*) is reported for any sequence with coverage below 0.9. Additional alerts can be reported at this stage (Table 3) based on unexpected characteristics of the set of returned alignments. For example, the *indf-strn* alert is reported if at least one alignment with a score of 25 bits or more occurs on each strand.

#### Alignment and feature mapping stage

Each complete sequence *S* is aligned to the covariance model for *M* (*S*) using Infernal’s *cmalign* program generating a Stockholm formatted alignment file. The alignment is then parsed to determine a mapping between sequence positions and model positions, from which per-feature coordinates for *S* are derived. The *cmalign* output Stockholm alignment file includes posterior probability values that indicate a level of confidence in each aligned nucleotide. These posterior probability values are used to identify features for which the endpoints are not confidently assigned by the model, which are reported as fatal alerts (*indf5loc* and *indf3loc*). Different fatal alerts are reported if an input sequence has a gap at a feature endpoint (*indf5gap* or *indf3loc*). Additional alerts are potentially reported at this stage as well (Table 3).

Standard CM alignment has high computational complexity in time and memory (*O*(*N*^4^) in time and *O*(*N* ^3^) in memory [20, 21]). Infernal utilizes a constrained alignment technique using bands derived from a first-pass HMM alignment of each sequence to make alignment practical [22, 23]. Input sequences that are dissimilar from the model’s RefSeq will have loose constraints, and the required memory for alignment may exceed 8Gb, in which case *cmalign* will exit in error without aligning the sequence. *v-annotate.pl* detects for which sequences this occurs, and reports an always fatal *unexdivg* alert.

#### Protein validation of CDS features

For each input sequence *S*, each predicted CDS feature of 30 or more nucleotides, as well as the full sequence *S*, is then used as a *blastx* query against the BLAST database of the RefSeq protein sequences created by *v-build.pl* for model *M* (*S*). The top *blastx* match for each RefSeq protein is compared to the CM-based prediction and alerts are generated if specific differences exist. For example, if the endpoints differ by more than five nucleotides on either the 5’ or 3’ ends, a *indf5pst* or *indf3pst* alert is reported. The main purpose of this stage is to identify any frameshift mutations in the input sequence, which may not have triggered any upstream alerts. Also at this stage, *insertnp* or *deletinp* alerts are reported for sequences with in-frame insertions or deletions longer than an indexer-specified taxon-specific threshold (set at 27 nucleotides by default), so that an indexer can check whether such a large insertion or deletion is plausible.

## Results and discussion

The VADR software package defines in one place all the checks that GenBank indexers want to do on the nucleotide sequences of incoming virus submissions. We define cautious operational semantics, summarized in Table 3, such that any sequence that might have a problem fails, so that an indexer can check the sequence.

### Obtaining norovirus and dengue virus sequences

To test the joint hypothesis that the VADR operational semantics are more cautious and more rigorous than what was done before its development (pre-2018), we collected a set of test sequences that were last modified no later than December 31, 2017. Specifically, we ran each of the following two Entrez Nucleotide queries on June 24, 2019:

1. Norovirus NOT chimeric AND 1989:2017[MDAT] AND 50:20000[slen]
2. Dengue NOT chimeric AND 1989:2017[MDAT] AND 50:20000[slen]

Distinct “complete genome” and “partial” sequence datasets were created to distinguish behavior on likely complete genomes from behavior on likely partial sequences. To separate the likely complete genomes, we used the empirical cumulative distribution functions of how many sequences returned by the above queries have each length, x, and are (not) annotated as “complete genome”. The selected thresholds were > 7380nt and > 10, 371nt for norovirus and dengue virus, respectively because almost all sequences labeled “complete genome” exceed these lengths. However, there were 5 norovirus and 27 dengue virus sequences shorter than these threshold lengths returned by the above Entrez queries that are annotated “complete genome”. Therefore, we revised the queries partitioning each sets of lengths into two disjoint intervals by using the “[slen]” attribute. We refer to the four datasets or selected subsets thereof as “NP”, “NC”, “DP”, and “DC” for Norovirus-Partial, Norovirus-Complete, Dengue-Partial and Dengue-Complete.

1. Norovirus NOT chimeric AND 1989:2017[MDAT] AND 50:7380[slen]
2. Norovirus NOT chimeric AND 1989:2017[MDAT] AND 7381:20000[slen]
3. Dengue NOT chimeric AND 1989:2017[MDAT] AND 50:10371[slen]
4. Dengue NOT chimeric AND 1989:2017[MDAT] AND 10372:20000[slen]

Queries 1 and 3 retrieved lists of 32,190 and 20,973 sequences respectively. Queries 2 and 4 retrieved lists of 1384 and 4580 sequences respectively. The partial sequences far outnumber the complete sequences for these two taxa. For this reason, we avoided design decisions in VADR that would rely on any specific part of the genome being present.

The sets of sequences returned by these queries could shrink over time if sequences are modified in the future. Two choices in our queries may be unexpected and deserve further explanation. First, we put the upper bound “2017” on the date because in 2018 we began using VADR to screen incoming sequences, so it would be unfair to include these more recent sequences in our testing here. In 2018, the versions of VADR were changing rapidly, so testing results from 2018 are omitted in the sequence counts at the beginning of Results, for which we started counting in January 2019. Second, we did not require that the strings Norovirus and Dengue be in the organism field, so as to include some sequences that could plausibly be misclassified taxonomically, and thereby test the behavior of VADR’s classification module. For simplicity and except where evaluating classification errors, we refer to the sequences as “norovirus sequences” and “dengue virus sequences”, even though some of them are known to belong to other taxa.

### VADR results for norovirus and dengue virus sequences

We ran the VADR v1.0 *v-annotate.pl* script with default parameters on the four sequence datasets. The numbers of sequences that pass and fail VADR from each of the four sets are shown in Table 4. For norovirus, about 92% of partial sequences and about 84% of complete genomes passed VADR. For dengue virus, about 82% of partial sequences and about 91% of complete genome sequences passed. The differences between the VADR pass rates on partial and complete sequences are partly due to idiosyncracies of typical primer pairs used for norovirus and dengue virus sequences and whether they capture portions of the genome that are easier (norovirus) or harder (dengue virus) to align.

**Table 4.** Summary of pass/fail outcomes for VADR on the full datasets

| dataset | VADR |  |
| --- | --- | --- |
|  | #pass/#fail | [pass fraction] |
| NC | 1157/227 | [0.836] |
| DC | 4171/409 | [0.911] |
| NP | 29488/2702 | [0.916] |
| DP | 17276/3697 | [0.824] |

Table 5 lists each type of fatal VADR alert observed in one of the four datasets with counts of instances reported and sequences for which one or more instances was reported. The most common alert, *peptrans*, occurs 6613 times in 1781 of the 59,127 sequences, approximately 3%. This alert does not indicate a unique problem itself, but rather is reported for mature peptides for which the parent CDS that is cleaved to form the mature peptide has a fatal alert, so it is redundant with at least one other alert. The next most common alert, *noannotn*, occurs for 2753 sequences, 2236 of which are in the DP dataset, indicating that no similar RefSeq was found for these sequences during the classification stage. Other alerts with more than 1000 instances include *indf5pst* and *indf3pst* which occur when the *blastx* protein-based alignment of a predicted CDS translation in the validation stage does not extend to with 5 nucleotides of the 5’ or 3’ ends of the nucleotide-based alignment. Thirty additional fatal alerts occurred for at least one sequence. Four fatal alert types did not occur for any sequence: *unexdivg* would be reported in the rare case that a sequence was recognized as similar to a RefSeq but too divergent to align within memory requirements; *lowsimis* would be reported if a dissimilar region occured outside all predicted features, which is unlikely for norovirus and dengue virus which have features nearly along their full length; *indfstrp* would be reported if *blastx* reported similarity to a CDS region on the negative strand, when nucleotide similarity is primarily recognized on the positive strand, and *incsbgrp* would be reported if a sequence is recognized as not belonging to a specified subgroup (e.g. Norovirus genotype), but was never reported in these tests because we did not specify subgroups. The *incsbgrp* alert was added to fit the design of the NCBI submission interface for norovirus in which submitters are asked to specify the genogroup, and it is triggered only when VADR judges that the sequence belongs to a different genogroup.

**Table 5.**
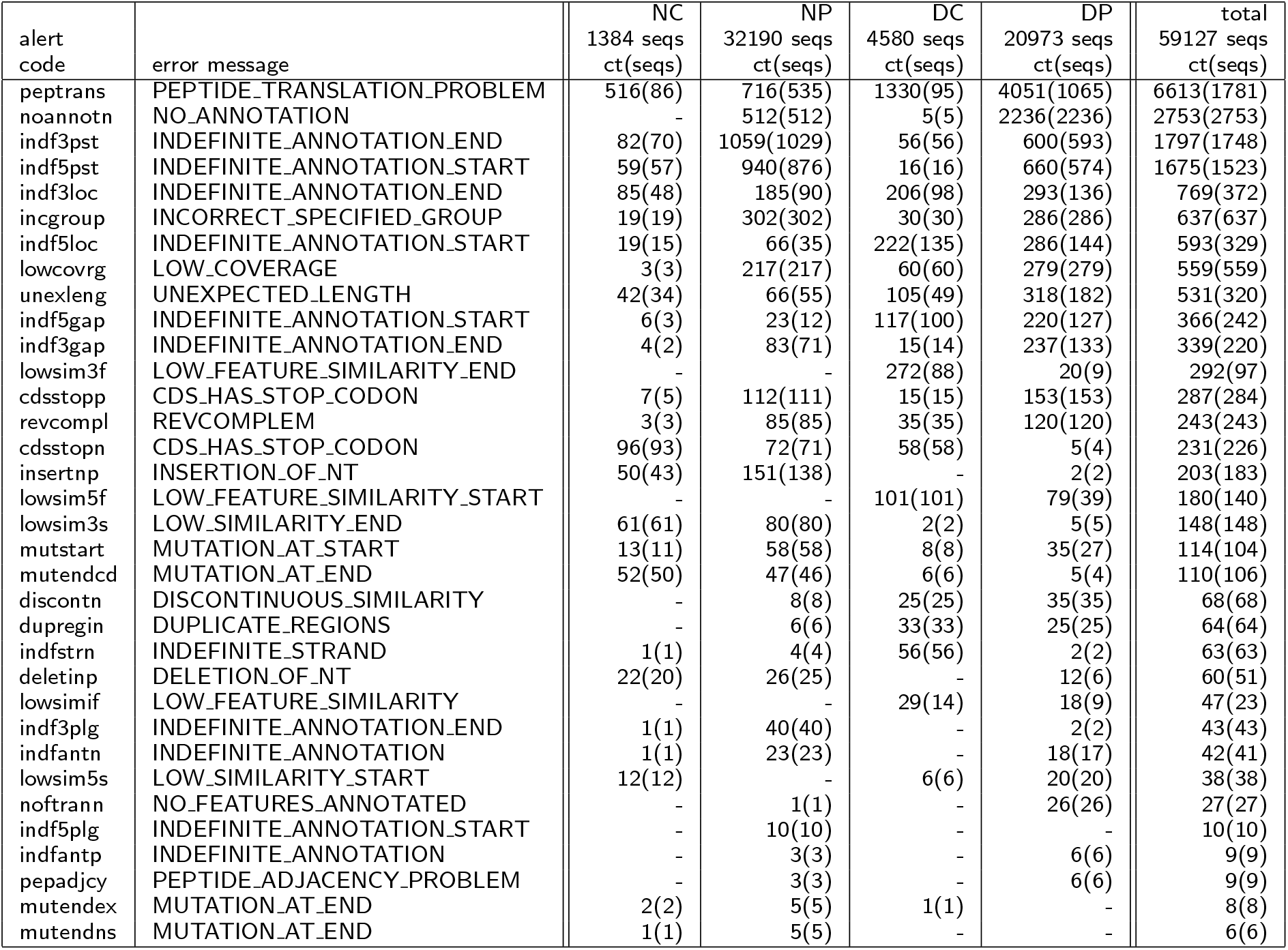
Counts of fatal VADR alerts reported for the test datasets. The 34 fatal alert codes reported at least once for any test dataset are listed sorted by total number of reports. 4 fatal alert types (unexdivg, lowsimis, incsbgrp and indfstrp.) were not reported for any of the four test sets and are not shown.See Table 3 for more information on alerts.

### Test datasets for comparison with VAPiD and VIGOR

Showing that VADR catches problems not caught before is not sufficient to prove that VADR is necessary to check virus submissions to GenBank, as existing software packages may be able to identify these problems as well. We compared the performance of VADR with that of VAPiD and VIGOR on randomly chosen sequences from the sets described above. From each of the four sets of sequences we randomly selected 200 to create four datasets of 200 sequences each for testing, with the condition that any sequence with 10 or more consecutive Ns, or with more than 50% ambiguous nucleotides was rejected from our sets of 200 because those sequences are flagged for manual review by NCBI indexers upon submission, outside the operation of VADR.

VAPiD expects complete genome sequences [7] and we verified that it reports spurious errors on the majority of partial sequences. Therefore, here, we report VAPiD tests only on the NC and DC (complete) datasets. The version of VIGOR we tested (VIGOR3) lacks a Dengue virus database, so we tested VIGOR only on the Norovirus NC and NP. Neither VAPiD nor VIGOR is currently designed to handle partial dengue virus sequences and this gives a simple justification for why VADR is needed to check dengue virus submissions.

### Defining pass/fail criteria for VAPiD and VIGOR

VAPiD and VIGOR were designed optimistically to prepare submissions to GenBank, while VADR was designed more pessimistically to identify unexpected characteristics in submitted sequences. VAPiD and VIGOR do catch some errors, but neither package has an overall definition of a sequence passing or failing as VADR does. Therefore, as described below, we defined operational semantics of pass/fail in VIGOR and VAPiD to make them more comparable to VADR.

To understand the operational semantics we imposed on VAPiD and VIGOR, it is necessary to begin with the point that they were designed to search the submitted sequence against one or more large databases of known sequences. These databases play a role analogous to the RefSeqs for VADR in that the best match in the database to the input sequence defines what features should be annotated. Because our tests used real sequences already in GenBank and the VAPiD and VIGOR databases also contain sequences already in GenBank, there turns out to be a non-negligible probability that our query sequences are already “known to” VAPiD and/or VIGOR.

For VAPiD, a sequence was considered to fail if: no reference matching it was found in the VAPiD database, the sequence was determined to be reverse complemented relative to the selected reference, or one or more “SEQ_FEAT” errors was listed in the output *errorsummary.val* file; all other sequences were considered to pass. Errors observed in our testing are “SEQ_FEAT.CdTransFail”, “SEQ_FEAT.NoStop”, “SEQ_FEAT.TransLen”, “SEQ_FEAT.StartCodon”, and “SEQ_FEAT.InternalStop”.

For VIGOR, any sequence that has no features annotated in the output *.tbl* file or for which one or more lines in the output *.at* file contained a code other than “OK”, “T5” or “T3” in the seventh field was considered to fail and all other sequences were considered to pass. Failure codes observed in our testing are “ES” and “FS” for detected embedded stops and detected frameshifts, respectively.

### Results of VADR, VAPiD, and VIGOR on test sets

Table 6 shows the number of sequences that pass and fail with each of the three methods for the datasets on which they were tested. Table 7 further compares the results of VADR and VAPiD on the NC and DC datasets, and Table 8 further compares the results of VADR and VIGOR on the NC and NP datasets. Because VADR is supposed to identify sequences with unusual properties that should be examined by an expert NCBI indexer before deposition into GenBank, the main goal of the comparison with VAPiD and VIGOR was to ensure that VADR identifies all the problems identified by the other two programs, and possibly more. To determine whether this is the case or not in our test sets, we examine the set of sequences that fail each method in turn below.

**Table 6.** Summary of pass/fail outcomes for VADR, VAPiD and VIGOR on the 200 sequence test datasets

| dataset | VADR<br>pass/fail | VAPiD<br>pass/fail | VIGOR<br>pass/fail |
| --- | --- | --- | --- |
| NC | 167/33 | 161/39 | 198/2 |
| DC | 189/11 | 196/4 | - |
| NP | 191/9 | - | 195/5 |
| DP | 163/37 | - | - |

**Table 7.** Comparison of pass/fail outcomes for VADR and VAPiD on the 200 sequence Norovirus-Complete (NC) and Dengue-Complete (DC) test datasets.

| dataset | Both<br>pass | Both<br>fail | VADR-pass<br>VAPiD-fail | VADR-fail<br>VAPiD-pass |
| --- | --- | --- | --- | --- |
| NC | 137 | 9 | 30 | 24 |
| DC | 188 | 3 | 1 | 8 |

**Table 8.** Comparison of pass/fail outcomes for VADR and VIGOR on the 200 sequence Norovirus-Complete (NC) and Norovirus-Partial (NP) test datasets.

| dataset | Both<br>pass | Both<br>fail | VADR-pass<br>VIGOR-fail | VADR-fail<br>VIGOR-pass |
| --- | --- | --- | --- | --- |
| NC | 167 | 2 | 0 | 31 |
| NP | 191 | 5 | 0 | 4 |

#### Test sequences that failed VAPiD

In our tests of the NC and DC datasets, VAPiD failed 43 total sequences; 12 of these were also failed by VADR and 31 were passed by VADR and VIGOR (Table 7). Of the 12, nine had an internal stop (SEQ_FEAT.InternalStop) or another CDS translation problem (SEQ_FEAT.CdsTransFail), one had a start codon problem (SEQ_FEAT.StartCodon), one was reverse complemented (FV536857.1; this was the only sequence of the 12 that failed VIGOR as well), and one failed because no similar reference was found for it. All 31 sequences that failed VAPiD but did not fail the other two programs (30 from the NC set and 1 from the DC set) were either 5’ truncated in the first CDS (Norovirus nonstructural polyprotein or Dengue virus polyprotein) or 3’ truncated in the final CDS (Norovirus VP2) or both, according to the VADR and VIGOR results, but were otherwise valid. These all failed VAPiD with at least one of the SEQ_FEAT.NoStop or SEQ_FEAT.CdTransFail errors, which is not surprising because VAPiD was designed for complete genomes and so does not expect truncated CDS sequences. Thus, we conclude that VADR successfully catches all the VAPiD-reported errors that should be caught.

#### Test sequences that failed VIGOR

In our tests of the NC and NP datasets, VIGOR failed seven total sequences, all seven of which also fail VADR (Table 8). Of the seven, three had premature stops, two were reverse complemented (one of these is FV536857.1), one had a frameshift and one failed because no similar reference was found for it. Thus, we conclude that VADR successfully catches all the VIGOR-reported errors that should be caught.

#### Test sequences that failed VADR

In our tests of the NC, NP, and DC datasets, VADR failed 53 sequences; 18 of these were also failed by VAPiD and/or VIGOR as mentioned above (exactly one sequence, FV536857.1, in the NC dataset failed all three methods), and 35 passed VAPiD and/or VIGOR (Table 8). These 35 sequences had issues that were not flagged by either of the other two programs but that indexers would like to manually review before possibly accepting the sequences into GenBank. Those issues include: 16 sequences with early stop codons compared with the closest RefSeq (*cdsstopn* alerts), 12 sequences for which a *blastx* alignment in the protein validation stage did not extend close enough to the 3’ boundary predicted from the nucleotide-based alignment (*indf5pst* or *indf3pst* alerts), ten sequences with low similarity to the RefSeq at the 5’ or 3’ end of the sequence or an annotated feature (*lowsim5f*, *lowsim3f*, or *lowsim3s* alerts), seven sequences where the 5’ or 3’ boundary of a feature was not aligned with sufficient confidence (*indf5loc*, *indf5gap*, or *indf3loc* alerts), five sequences that were expected to be Norovirus but were classified as Sapovirus, another *Caliciviridae* genus (*incgroup* alerts), five sequences with too large of an insertion or deletion in a *blastx* alignment (*insertnp* or *deletinp* alerts), and one sequence not recognized by any of the *Caliciviridae* or *Flaviviridae* models (a Salivirus from the *Picornaviridae* family, *noannotn* alert). Twelve of the 35 sequences had more than one of the above listed alerts. For the sequences with truncated *blastx* alignments, we observed that identical or similar truncated alignments were found in VIGOR, but in the current design of VIGOR, truncated nucleotide-to-protein alignments do not trigger an error, at least in these cases.

In the DP dataset, for which only VADR was tested, 37 sequences failed. Twenty-eight failed because no similarity to any RefSeq in the reference library was detected (*noannotn* alert). Twenty of these are likely not dengue virus sequences but rather from *Aedes aergypti* or *Aedes albopictus*, the mosquito vectors of Dengue virus as they contain one of those species names in the sequence description. Three of the other nine sequences that failed did so because they are reverse complemented (*revcompl* alert). Three others fail because the *blastx* alignments do not extend far enough relative to the nucleotide-based CDS endpoints (*indf5pst* or *indf3pst* alerts). One failed because it had an internal stop in the polyprotein CDS (*cdsstopp* alert). One failed because its stem loop feature that is supposed to be found upstream of the CDS lacked similarity to the RefSeq in the first 30 nucleotides (*lowsim5f* alert). The final sequence failed because it was classified as coming from a different species in family *Flaviviridae*, namely West Nile virus (*incgroup* alert).

#### Definition of false positive and false negative rates for the NC sequences

For the NC data set, which was tested with all three packages, we attempted to make plausible definitions of false positives (FPs) and false negatives (FNs), so as to compare the three programs by standard measures in statistics. The sequences that are not from the Norovirus taxon are useful tests for quality control, but are implausible submissions, so we exclude the 5 sequences that are not classified as Norovirus in GenBank. Additionally, we exclude the 45 sequences in the NC set that VADR determines are truncated on the 5’ or 3’ end (that either begin 3’ of the start of the nonstructural polyprotein CDS or end 5’ of the end of the VP2 CDS) because VAPiD is not designed to handle incomplete sequences. This leaves 150 of the original 200 sequences in the NC set for defining FP and FN rates. We assert that the 129 sequences that pass all three programs are all positives, which is reasonable for our specific purpose of comparing only these three programs.

We manually examined the remaining 21 sequences and determined that 3 of them have undeniable serious errors and so should be classified as ‘negatives’. FV536857.1, which matches best to a non-norovirus RefSeq in VADR’s model library (NC 006875) and appears to be reverse complemented, is failed by all three programs and so is counted for all three as a true negative (TN). FJ446720.1, which has an early in-frame stop codon 690 nt 5’ of the expected stop site, is appropriately failed by VADR and VIGOR (TN) but passed by VAPiD (FP). KY905331.1 lacks a valid start codon in the nonstructural polyprotein CDS and is failed by VADR and VAPiD (TN) but passed by VIGOR (FP). We cautiously defined the remaining 18 sequences, all of which are failed by VADR but passed by VIGOR and VAPiD, as positives because each sequence is arguably valid but is just too different from the matching RefSeq model to pass VADR. These 18 sequences are TPs for VAPiD and VIGOR and FNs for VADR.

Based on this analysis for the NC results, VADR had a false positive rate of 0 and a false negative rate of 0.12 (18/147). Both VAPiD and VIGOR had a false positive rate of 0.33 (1/3) and a false negative rate of 0. Because one of the design goals of VADR is to prevent problematic sequences from automatic deposition into GenBank, it is more important to minimize the FP rate than the FN rate. While VADR’s FN rate is considerably higher than that of VIGOR and VAPiD, the 18 FN sequences that fail VADR all deviate from the nearest RefSeq in specific ways such that GenBank indexers want to manually examine them. Later, we calculate FP and FN rates for new norovirus and dengue submissions evaluated with VADR which show that the FN rate can be significantly lower in practice.

In summary, to allow for automatic processing of high quality virus sequences, VADR is designed to identify when sequences deviate from the closest RefSeq in various ways and construct informative alerts and output messages about those deviations. VADR catches all the problems that VAPiD and VIGOR catch and numerous other problems. Because of VADR’s usage to check submissions, we can tolerate some false negatives, but the false positive rate should be as close to 0 as possible.

#### Relevance of reference database size

The three programs differ in the size of the reference databases they use to map annotation onto input sequences. VADR’s reference library includes nine norovirus RefSeqs and four dengue virus RefSeqs. VAPiD’s reference library contains more than 800 norovirus and more than 4000 dengue virus sequences. VIGOR’s library contains more than 200 total norovirus proteins (polyprotein, VP1 and VP2) and more than 50 norovirus mature peptides. Because of VADR’s role in controlling which sequences are automatically entered into GenBank and how those sequences are annotated, having trustworthy and consistent annotations in the reference library is crucial. We chose RefSeqs as the basis for our library for this reason, after reviewing, updating and in some cases creating new RefSeq records with the intention that they would serve as references to the wider research community beyond the scope of VADR-based annotation.

Larger libraries are more likely to contain mistakes or anomalies that should not be mapped onto incoming sequences that may be automaticlly deposited in GenBank. An example is the sequence FJ446720.1 from the NC dataset, which is in the VAPiD reference library but lacks annotation of the VP1 CDS. Both VADR and VIGOR failed this sequence due to a premature stop in the putative VP1 CDS, but it passed VAPiD because its reference sequence (itself) lacks VP1 annotation and so the VP1 region is not examined for premature stops. This sequence may be biologically valid, but should be reviewed by GenBank indexers as opposed to being automatically deposited into GenBank.

On the other hand, a larger reference database can contain more diversity than a smaller one, and the most common VADR failure in the NC dataset is due to an early stop codon in the nonstructural polyprotein CDS by three nucleotides (11 of the 16 *cdsstopn* alerts in the set of 35 sequences mentioned above). This failure may have been avoided with a larger reference database that included a norovirus sequence with this three nucleotide shorter CDS variant (an example is AB933745.1). We plan to add to VADR’s reference library as we find areas of sequence space that it does not adequately cover; for example, while the manuscript was out for review, 10 norovirus RefSeqs were added for internal testing. We also plan to extend VADR to use profiles built from multiple alignments instead of single sequences which should enhance its ability to analyze and annotate some sequences that are divergent from available RefSeqs.

#### Agreement on passing sequences

The above discussion focuses on failing sequences, and does not address the validity of passing sequences or the correctness of their annotation. In our testing of VADR since May 2018, indexers have manually reviewed the sequences that pass VADR and have largely agreed that they should indeed pass. Where they have disagreed, we have modified VADR during its development to fail the sequences in question. Additionally, in our testing of the NP and NC datasets, when VADR and VIGOR both pass a sequence they nearly always have the same annotations: 703 of the 705 (99.7%) of the CDS annotations with consistent product names (VP1, VP2 or nonstructural polyprotein) had identical coordinates in the VADR and VIGOR output.

### Sequences already evaluated with VADR

To date, VADR has been used to evaluate hundreds of norovirus and dengue virus sequences submitted to GenBank. During intermediate phases of testing, we recorded in detail the fate of every submitted sequence into six categories. Among 922 norovirus sequences submitted in 2019 during this phase, 898 (97%) were accepted automatically, 8 were accepted by an indexer without changes, 8 were accepted by an indexer after making changes, and 8 were sent back to the submitter. Among 2644 dengue virus sequences submitted from the beginning of 2019 until intermediate testing stopped on February 14, 2020, 2520 (95%) were accepted automatically, 15 were accepted by an indexer without changes, 96 were accepted by an indexer after making changes, 9 were sent back to the submitter, and 4 had a more complicated fate. After VADR was put into full production usage for norovirus, the record-keeping was reduced to record only how many sequences are evaluated by VADR and how many sequences pass or fail. Among the first 3143 norovirus sequences evaluated by VADR in production mode, 2809 (89%) passed while 334 (11%) failed. If these data are representative of norovirus and dengue virus submissions, the expected reduction in the number of sequences that need to be manually reviewed by NCBI indexers (which prior to development of VADR was 100%) will be about 10-fold for norovirus submissions and about 20-fold for dengue virus submissions.

In the context of the new submissions, there is virtually no risk that a sequence of the wrong taxon would be submitted via the new GenBank submission portal interfaces for norovirus submissions and dengue virus submission. Therefore, we define that a sequences is a *false positive* if it passes VADR, but a GenBank indexer deems using other tools that the sequence has an error. Similarly, a sequence is a *false negative* if it fails VADR, but a GenBank indexer deems that the sequence should pass with no errors. Using these definitions and the counts in the previous paragraph, VADR’s false positive rates are 0 for both norovirus and dengue virus. The false negative rates are 0.009 (8/906) for norovirus and 0.006 (15/2535) for dengue virus.

### VADR-based annotation of mature peptides and structural RNA features

The central problems in checking and annotating viral genomes and bacterial genomes differ [2]. Bacterial genomes are typically larger with hundreds of genes, many of which may be uncharacterized. Hence, assembly and gene prediction are two of the difficult problems in bacterial annotation. Virus genomes are typically smaller with only a handful of genes, but the biological phenomena of cleavage of proteins into mature peptides [24] and ribosomal slippage [25, 26, 27] are prominent. Surprisingly, while 97% (31,086 of 32,190) and 85% (17,900 of 20,973) of non-RefSeq norovirus and dengue virus sequences, respectively, last modified between January 1, 1989 and December 31, 2017, had at least one CDS annotated, only 1.4% (443 of 31,086) and 6.4% (1,157 of 17,900) of those those had any mature peptides annotated, even though mature peptide cleavage occurs in all viruses of the corresponding genera. VADR fixes this omission; all norovirus and dengue virus sequences in GenBank annotated with VADR have mature peptides predicted, where possible.

As noted above, VAPiD and VIGOR do not attempt to annotate features other than coding sequences: both annotate CDS, and VIGOR annotates mat peptides. VADR has the added capability of annotating any sequence feature that is also annotated in the RefSeq, including conserved structural RNA elements which, though present in many viral genomes [28, 29], are typically not annotated in GenBank. We added annotation of an ncRNA feature, a subgenomic flavivirus RNA (sfRNA), and associated stem loop features, to the four dengue virus RefSeqs in preparation of VADR use for dengue sequence submissions. These structural RNA elements are relevant in pathogenicity and evasion of the host immune system in at least some flaviviruses [13, 30, 31]. Incoming dengue virus sequence submissions will now include these RNA annotations because of VADR, which employs covariance models of both the conserved sequence and secondary structure of the RNA elements. In the set of 4171 sequences that passed VADR in the full DC dataset of 4580 sequences, VADR annotated between 4 and 9 stem loop features and exactly 1 ncRNA feature in each sequence for a total of 35,676 stem loop features and 4171 ncRNA features. In the set of 17,276 sequences that passed VADR in the full 20,973 sequence DP dataset, VADR annotated between 1 and 6 stem loop features in 2335 sequences, and exactly one ncRNA feature in 623 of those 2327 sequences for a total of 5157 stem loop features and 623 ncRNA features.

VADR can also annotate CDS that involve ribosomal slippage and that capability will become evident when it begins to be used for Hepacivirus, West Nile virus, or ebolaviruses. VADR’s full sequence alignment-based annotation strategy enables annotation of any feature that has nucleotide positional information in the RefSeq annotation. This includes discontiguous features, including multi-segment genes and features as short as a single nucleotide.

### Limitations and future directions

One obvious limitation of VADR at present is that it is used at NCBI only for norovirus and dengue virus sequences, but not yet for other viruses. Near-term plans include to expand the usage to many other members of the family *Flaviviridae*, including Hepacivirus C, West Nile virus and Zika virus. VADR was not designed for viruses with genomes larger than 25Kb such as eukaryotic DNA viruses such as Epstein-Barr virus (~170Kb). For these larger viruses, VADR would require an impractical amount of memory and time due to the high complexity of the CM algorithms. In future work, we hope to develop new methods to alleviate this restriction. Due to the COVID-19 outbreak beginning in December 2019, we have made 55 *Coronaviridae* models available for download and analysis of coronavirus sequences including SARS-CoV-2, even though VADR was not designed for sequences as large as coronavirus genomes (~30Kb). It is recommended that at least 64Gb of RAM are available when analyzing full length coronavirus genomes. Download and usage instructions for these models can be found on github (https://github.com/nawrockie/vadr/wiki/Available-VADR-model-files).

Users may construct VADR models for any viral species represented by a RefSeq or other GenBank sequence up to 25Kb, but it is important to make sure that all the features for the sequence are properly annotated with the current nomenclature. Users may write to either to inquire as to which viral RefSeqs have been vetted to the standards needed in VADR and required by GenBank indexers, or to suggest corrections or request updates to RefSeq annotations.

Another limitation of the current usage of VADR is that each model is built from a single RefSeq sequence. Valid sequences that are sufficiently dissimilar from all RefSeq-based models will fail because alerts designed to identify dissimilar sequences will be reported. The failures of some sequences from the test datasets that pass VAPiD and/or VIGOR can be attributed to this, as discussed in the Results section. During testing with incoming norovirus sequence submissions, three additional RefSeqs, included in the nine norovirus RefSeqs used for the tests described herein, were created to address this issue. Stimulated by some recent norovirus submissions with sequences dissimilar from the nine RefSeqs in the VADR 1.0 library and by a newly published norovirus genotyping paradigm [32] that suggested additional candidate RefSeqs more similar to the new sequences, we selected and curated 11 additional norovirus RefSeqs to reach a total of 20. Ten of the additional 11 RefSeqs are being tested on norovirus submissions since early 2020. VADR can be run on such sequences with 19 or 20 models for norovirus instead of nine models. Whichever ones of the additional 11 norovirus RefSeqs prove useful in internal testing will have their models included in a future released version of VADR and be used in automated processing of submissions.

An alternative strategy to building models from a single RefSeq is to create and use profile models (profile HMMs and CMs) from trusted sequence alignments of multiple representative sequences that cover the known diversity of the virus species or subspecies being modelled. Profile-based methods are more sensitive at homology detection [33, 34, 35, 36] then single sequence based methods and so this strategy may improve performance. Extending VADR to profiles was envisioned from its design inception and the *cmbuild* program which creates VADR’s CM files can take as input a multiple alignment. All sequences in the alignment the profile is constructed from should include the same set of features with start and end points aligned, which will limit the phylogenetic breadth of some alignments. For example, among noroviruses, eight of the nine RefSeqs encode three proteins, but murine norovirus (represented by NC 08311) encodes four proteins. A similar problem arises among different subtypes of West Nile virus that may or may not encode WARF4 [37].

VADR is also flexible as to what types of sequences it models, and it is not restricted to viruses. Any conserved sequence feature, such as a structural RNA gene or region, or part or all of a viroid, can be modelled by VADR. The accuracy of VADR annotations for any sequence depends on how similar it is to the reference sequence(s) from which the model was built.

Although VADR uses Infernal, a program capable of modelling RNA secondary structure, its models will not include structure information unless the user provides it as input to *v-build.pl*. For models without secondary structure it could be reasonably argued that VADR should use profile HMMs instead of CMs for the alignment stage, because HMMs are more efficient to compute with than CMs and in the absence of secondary structure are essentially equivalent models. However, we opted to always use CMs for the alignment stage because the popular HMM package HMMER3 is not well-suited to accurately defining alignment endpoints, as it is restricted to local alignment only, whereas Infernal allows global alignment with respect to the sequence, and consequently can identify at least some boundaries more accurately.

For some potential users, the requirement to use VADR on either Linux or Mac OS X via the command line is a limitation. Therefore, planning is underway to develop a Dockerized version of VADR that could be used via cloud computing.

The relatively slow speed of VADR is also a limitation. Running as a single execution threads on a 2.93 GHz Intel Xeon processor, the *v-build.pl* script takes about 15 minutes for a typical calicivirus genome, and about 25 minutes for a typical flavivirus genome, but this step needs to only be run once per RefSeq. The *v-annotate.pl* script takes about 1.5 seconds or 30 seconds for partial or full-length Norovirus sequences, respectively, and about 9 seconds or 90 seconds for partial or full-length Dengue sequences, respectively. These speeds are about 8 times (NC dataset) or 20 times (DC dataset) slower than VAPiD, and 6 times (NC dataset) or 1.5 times (NP dataset) slower than VIGOR. The two most expensive stages of *v-annotate.pl* are the classification and alignment stages. To speed up sequence processing, the *v-annotate.pl* script can be run in parallel without impacting the results, by partitioning the input sequence file, running each partition separately in parallel, and then concatenating the output files. Additionally, the *-p* option allows users with access to a compute cluster to do this partitioning and parallelization automatically.

## Conclusions

VADR is NCBI’s new production software system for checking and annotating non-influenza virus sequence submissions to GenBank. VADR handles both whole genomes and partial genomes. From January 1, 2019 through October 8, 2019, VADR was used to check and annotate 4065 norovirus sequences and 1702 dengue virus sequences. Norovirus and dengue virus researchers can scrutinize the newly annotated sequences and install and use VADR to check sequences that they generate in the future before submitting them to GenBank. VADR implements rigorously documented operational semantics for characterizing sequence problems and other unexpected characteristics, which should simplify the interaction between sequence submitters and GenBank indexers when submitted sequences fail. VADR will be gradually adapted to include more of the viruses in Table 1 and their relatives starting with other members of the family *Flaviviridae*.

## Supporting information

Supplemental material tarball

## Availability and requirements

Project name: VADR

Project home page: https://github.com/nawrockie/vadr

Operating system(s): Linux, Mac OS X

Programming language: Perl

Other requirements: Bio-Easel v0.09, BLAST+ v2.9.0, Infernal v1.1.3, Sequip v0.03 (see Table 2)

License: public domain

Any restrictions to use by non-academics: none

## Abbreviations

NCBI: National Center for Biotechnology Information
VADR: Viral Annotation DefineR
BLAST: Basic Local Alignment Search Tool
CM: Covariance Model
HMM: Hidden Markov Model
nt: nucleotides
Kb: kilobase (1000 nucleotides)
ncRNA: noncoding RNA
mat_peptide: mature peptide
NP: Norovirus Partial dataset
NC: Norovirus Complete dataset
DP: Dengue virus Partial dataset
DC: Dengue virus Complete dataset

## Ethics approval and consent to participate

Not applicable.

## Consent for publication

Not applicable.

## Availability of data and materials

All data generated or analyzed during this study are included in this published article, its supplementary material, or NCBI’s GenBank database. Code is available on github (https://github.com/nawrockie/vadr). The supplementary material includes instructions for reproducing the comparisons of VADR, VAPiD and VIGOR reported in the article.

## Author’s contributions

AAS wrote the protein validation module and EPN wrote the remainder of the code. ELH updated RefSeq annotation and developed test cases. LY tested the software on sequence submissions and developed test cases. LS wrote an early version of the classification module. JRB and IK-M helped develop the initial specifications for VADR and managed the work of ELH and LY, respectively. AAS and EPN conceived of and designed the software and wrote the paper. All authors helped in the development of the software and read and approved the final manuscript.

## Acknowledgements

We thank Estéban Finol and Mariano Garcia-Blanco for providing predicted RNA structures for the Dengue virus RefSeqs, Alex Kotliarov for incorporating VADR into the GenBank submission portal system, Ron Patterson and his team for managing NCBI’s computing resources, and Vince Calhoun for testing VADR and helpful feedback on the manuscript. This research was supported by the Intramural Research Program of the NIH, National Library of Medicine and National Cancer Institute.

## Additional Files

Additional file 1 — vadr-1.0-paper-supplementary-material.tar.gz

Sequence files, scripts, and instructions for reproducing Norovirus and Dengue virus tests reported in the article. This is a gzipped tarball that can be unpacked with the command ‘tar xf vadr-1.0-paper-supplementary-material.tar.gz’, and includes two README files with instructions on reproducing the tests and reported results.

