## Supplemental material tarball for "VADR: validation and annotation of virus sequence submissions to GenBank": README-processing-test-results.txt

EPN, Tue Nov  5 13:39:19 2019

README-processing-test-results.txt

Notes on processing the results from running VADR, VIGOR and VAPiD on
the test sequence datasets described in "VADR: validation and
annotation of virus sequence submissions to GenBank"

(Lines of this file that are commands are prefixed with '$')

-------------
To reproduce the data presented in the paper "VADR: validation and
annotation of virus sequence submissions to GenBank" that corresponds
to the runs of VADR, VAPID and VIGOR on the test sequence datasets,
perform the following steps 1-8.

The RELEVANCE TO PAPER section in each step below explains its
relevance to the data presented in the paper.

---------------------------------------------------------------------- 

1. Install and run the programs VADR, VAPiD and VIGOR on the test
   sequence datasets as explained in the file README-running-tests.txt

   RELEVANCE TO PAPER: runs the programs and creates output that is
   eventually processed to make tables and derive other statistics in
   the paper. 

---------------------------------------------------------------------- 

2. Set-up environment variables to be able to process the results by 
   first modifying the file 'scripts/set-environment.sh' 
   as necessary to point each environment variable to the appropriate
   path in your file system.

   Then run the following command:

$ source scripts/set-environment.sh

   This will set the following environment variables:
   VADRPAPER_SUPPDIR to where you unpacked the supplementary material tarball.
   VADRPAPER_VADRDIR to where the VADR test run output files are
   VADRPAPER_VAPIDDIR to where the VAPiD test run output files are
   VADRPAPER_VIGORDIR to where the VIGOR test run output files are
   VADRPAPER_SCRIPTSDIR to the scripts directory in the supplementary
                         material ($VADRPAPER_SUPPDIR/scripts)
 
   RELEVANCE TO PAPER: sets up environment variables so that steps 3-8
   below will run and create data for the paper.
   
---------------------------------------------------------------------- 

3. Create the summary files of the results from each of the three
   programs (this puts the results from the three programs into the
   same format to make processing and comparing them easier) using the 
   following command:

$ $VADRPAPER_SCRIPTSDIR/make-summaries.sh

   The 'make-summaries.sh' script calls the following perl scripts
   (also in $VADRPAPER_SCRIPTSDIR):
   vapid-out-and-errorsummary-to-seqsum.pl
   vigor-at-to-seqsum-and-ftrsum.pl
   vadr-sqa-and-ftr-to-seqsum-and-ftrsum.pl

   RELEVANCE TO PAPER: creates intermediate files that are eventually
   processed into the tables in the paper in step 5.

---------------------------------------------------------------------- 

4. Combine .seqsum files and output summary of number of pass/fail
   with the following command:

$ $VADRPAPER_SCRIPTSDIR/combine-summaries.sh > pass-fail-counts.txt

   The 'combine-summaries.sh' script calls the following perl script
   (also in $VADRPAPER_SCRIPTSDIR):
   combine-seqsums.pl

   RELEVANCE TO PAPER: creates intermediate files that are processed
   into the tables in the paper in step 5.

---------------------------------------------------------------------- 

5. Create latex tables for paper with the following commands:

$ $VADRPAPER_SCRIPTSDIR/combine-summaries-for-tbl.sh | perl $VADRPAPER_SCRIPTSDIR/vadr-make-results-tables.pl > tables-results.tex  
$ perl $VADRPAPER_SCRIPTSDIR/vadr-make-alertlist-table.pl $VADRPAPER_SUPPDIR/vadr.1p0.alt_list > table-alertlist.tex
$ perl $VADRPAPER_SCRIPTSDIR/vadr-make-alertcounts-table.pl $VADRPAPER_SUPPDIR/vadr.1p0.alt_list test > table-alertcounts.tex
$ perl $VADRPAPER_SCRIPTSDIR/vadr-make-full-results-table.pl > table-full-results.tex
$ perl $VADRPAPER_SCRIPTSDIR/vadr-make-alertcounts-table.pl $VADRPAPER_SUPPDIR/vadr.1p0.alt_list full > table-full-alertcounts.tex

   Note: The '$VADRPAPER_SUPPDIR/vadr.1p0.alt_list' file was created
   with the command:
   $ $VADRSCRIPTSDIR/v-annotate.pl --alt_list > vadr.1p0.alt_list.txt

   Versions of these .tex files used in the submitted version of the
   paper are in the supplementary data in the latex-tables/ subdirectory.

   RELEVANCE TO PAPER: 
   table-alertlist.tex         includes Table  3           from submitted paper.
   table-full-results.tex      includes Table  4           from submitted paper.
   table-full-alertcounts.tex  includes Table  5           from submitted paper.
   tables-results.tex          includes Tables 6, 7, and 8 from submitted paper.
   table-alertcounts.tex       includes a modified version of Table 5
                               that pertains to the 200 sequence
                               datasets only (not in paper)

---------------------------------------------------------------------- 

6. Compare VIGOR and VADR Norovirus CDS annotated sequence coordinates
   in sequences that pass both programs and have consistent product
   names with the command: 

$ $VADRPAPER_SCRIPTSDIR/vadr-vigor-compare-ftrsums.sh > vadr-vigor-compare-ftrsums.out

   The 'vadr-vigor-compare-ftrsums.sh' script calls the following perl
   script (also in $VADRPAPER_SCRIPTSDIR):
   compare-two-ftrsums.pl

   NOTE: compare-two-ftrsums.pl (called in
   vadr-vigor-compare-ftrsums.sh) REMOVES the string 'capsid_protein_'
   from the beginning of product names in the VADR and VIGOR ftrsum
   output files.

   NOTE: vigor-at-to-seqsum-and-ftrsum.pl REMOVES the strings
   'truncated ' and 'putative ' from the beginning, and ', fragment',
   'C-terminal' and 'N-terminal' from the end of CDS product names. So
   those have ALREADY BEEN REMOVED in the .ftrsum output files output
   output from vigor-at-to-seqsum-and-ftrsum.pl called by
   make-summaries.sh in step 2 above.

   RELEVANCE TO PAPER: the file vadr-vigor-compare-ftrsums.out
   includes the counts cited in the paper towards the end of the
   Results section: 
   "703 of the 705 (99.7%) of
   the CDS annotations with consistent product names
   (VP1, VP2 or nonstructural polyprotein) had identi-
   cal coordinates in the VADR and VIGOR output."

---------------------------------------------------------------------- 

7. Count VADR RNA annotations in full test set with the command:

$ $VADRPAPER_SCRIPTSDIR/vadr-count-rnas-full.sh > vadr-rnas.full.out

   RELEVANCE TO PAPER: the file vadr-rnas.full.out includes the counts
   of RNA annotations included in the second to last paragraph of the
   Results section.

---------------------------------------------------------------------- 

8. Count number of sequences with >= 1 CDS annotation and >=1 mature
   peptide annotation. These require that edirect be installed on your
   system and in your path (see
   https://www.ncbi.nlm.nih.gov/books/NBK179288/).

$ $VADRPAPER_SCRIPTSDIR/genbank-fetch-cds-and-mp.sh
$ $VADRPAPER_SCRIPTSDIR/genbank-count-cds-and-mp.sh > cds-and-mp-counts.txt
   
   The 'genbank-fetch-cds-and-mp.sh' script calls the following perl
   script (also in $VADRPAPER_SCRIPTSDIR):
   acclist2mp-and-cds.pl

   RELEVANCE TO PAPER: the file cds-and-mp-counts.txt includes the
   counts related to CDS and mat_peptide annotations in the Discussion
   section.

----------------------------------------
Questions, contact
----------------------------------------
