## Supplemental material tarball for "VADR: validation and annotation of virus sequence submissions to GenBank": README-running-tests.txt

EPN, Tue Nov  5 06:10:24 2019

README-running-tests.txt

Notes on installing and running VADR, VIGOR and VAPiD on the test
sequence datasets described in "VADR: validation and annotation of
virus sequence submissions to GenBank".

After the runs described in this file are performed, see 
README-processing-test-results.txt for information on how the data
presented in the paper was obtained from the output of the runs.

(Lines of this file that are commands are prefixed with '$')

Outline of this file:
CONSTRUCTING TEST DATASETS
INSTALLING AND RUNNING VADR
INSTALLING AND RUNNING VAPID
INSTALLING AND RUNNING VIGOR

==========================
CONSTRUCTING TEST DATASETS
==========================
The following Nucleotide Entrez queries
(https://www.ncbi.nlm.nih.gov/nuccore/) were used on June 24,2019, as
reported in the Results section of the paper:

1. Norovirus NOT chimeric AND 1989:2017[MDAT] AND 50:7380[slen]
2. Norovirus NOT chimeric AND 1989:2017[MDAT] AND 7381:20000[slen]
3. Dengue NOT chimeric AND 1989:2017[MDAT] AND 50:10371[slen]
4. Dengue NOT chimeric AND 1989:2017[MDAT] AND 10372:20000[slen]

Query 1 returns 32,190 accessions (Norovirus-Partial 'NP' dataset)
Query 2 returns  1,384 accessions (Norovirus-Complete 'NC' dataset)
Query 3 returns 20,973 accessions (Dengue-Partial 'DP' dataset)
Query 4 returns  4,580 accessions (Dengue-Complete 'DC' dataset)

From each dataset 200 random sequences were selected using the
'esl-selectn' program of Infernal 1.1.2, e.g.:

$ esl-selectn 200 np.list > np.r200.list

*At this step* sequences with too many ambiguous (any character
except, 'A', 'C', 'G' or 'T') characters were rejected and another
randomly selected sequence was chosen to replace it. A sequence has
"too many Ns" if it has either 10 or more consecutive Ns or is more
than 50% ambiguous characters. Such sequences were identified using
the 'count-ambigs.pl' and 'longest-N-stretch.pl' Perl scripts in the 
'scripts' subdirectory of the supplementary data. 
     
Relevant files in the supplementary data: 
-----------------------------------------
In seq-files/ subdir:

np.fa:        FASTA-formatted sequence file of full NP dataset
np.r200.fa:   FASTA-formatted sequence file of 200 seq NP dataset
np.list:      list of accessions in full NP dataset
np.r200.list: list of accessions in 200 seq NP dataset
nc.fa:        FASTA-formatted sequence file of full NC dataset
nc.r200.fa:   FASTA-formatted sequence file of 200 seq NC dataset
nc.list:      list of accessions in full NC dataset
nc.r200.list: list of accessions in 200 seq NC dataset
dp.fa:        FASTA-formatted sequence file of full DP dataset
dp.r200.fa:   FASTA-formatted sequence file of 200 seq DP dataset
dp.list:      list of accessions in full DP dataset
dp.r200.list: list of accessions in 200 seq DP dataset
dc.fa:        FASTA-formatted sequence file of full DC dataset
dc.r200.fa:   FASTA-formatted sequence file of 200 seq DC dataset
dc.list:      list of accessions in full DC dataset
dc.r200.list: list of accessions in 200 seq DC dataset

--
In seq-files/vapid-indi-files/ subdir:
(These files are used for VAPiD runs as explained more below)

For <i> in range 1..200:

nc.r200.fa.<i>         single sequence from nc.r200.fa 
nc.r200.<i>.csv        metadata file for VAPiD for this sequence 
nc.r200.<i>.namelist   name of sequence in nc.r200.fa.<i>

dc.r200.fa.<i>         single sequence from dc.r200.fa 
dc.r200.<i>.csv        metadata file for VAPiD for this sequedce 
dc.r200.<i>.namelist   name of sequence in dc.r200.fa.<i>

--
In scripts/ subdir:
count-ambigs.pl: script for counting ambiguous characters in
                 each sequence in a nucleotide fasta file
longest-N-stretch.pl: script for identifying longest stretch
                      of Ns in each sequence in a fasta file


-----------------------------------------

===========================
INSTALLING AND RUNNING VADR
===========================

VADR 1.0 was installed along with its dependencies using the
installation instructions in the README.md available here:

https://github.com/nawrockie/vadr/blob/1.0/README.md

VADR was run on the full set of sequences using a compute farm/cluster
with the following commands:

$ v-annotate.pl -p --group Norovirus -f np.fa va-np > va-np.out
$ v-annotate.pl -p --group Norovirus -f nc.fa va-nc > va-nc.out
$ v-annotate.pl -p --group Dengue -f dp.fa va-dp > va-dp.out
$ v-annotate.pl -p --group Dengue -f dc.fa va-dc > va-dc.out

VADR was run on the 200 sequence sets on a single processor with the
following commands:

$ v-annotate.pl --group Norovirus -f np.r200.fa va-np.r200 > va-np.r200.out
$ v-annotate.pl --group Norovirus -f nc.r200.fa va-nc.r200 > va-nc.r200.out
$ v-annotate.pl --group Dengue -f dp.r200.fa va-dp.r200 > va-dp.r200.out
$ v-annotate.pl --group Dengue -f dc.r200.fa va-dc.r200 > va-dc.r200.out

Results were processed as explained in
README-processing-test-results.txt

============================
INSTALLING AND RUNNING VAPID
============================

Installed VAPiD v1.6.3 by following installation instructions in
https://github.com/rcs333/VAPiD/blob/master/README.md
(commit 23299e2)

To get VAPiD to run in my hands I had to change the vapid.py file at
line 350 from :

                     allow_one = Falseå

To 
                     allow_one = False


$ python VAPiD-master/vapid.py np.r200.fa VAPiD-master/example.sbt --metadata_loc np.r200.csv

---
VAPiD was run individually on each sequence in the 200 sequence sets
on a single processor with a bash shell script like this:

for ((i = 1; i <= 200; i++); do 
  python vapid.py seq-files/vapid-indi-files/nc.r200.fa.$i VAPiD-master/example.sbt --metadata_loc seq-files/vapid-indi-files/nc.r200.$i.csv > vp-nc.r200.$i.out 2> vp-nc.r200.$i.err
  python vapid.py seq-files/vapid-indi-files/dc.r200.fa.$i VAPiD-master/example.sbt --metadata_loc seq-files/vapid-indi-files/dc.r200.$i.csv > vp-dc.r200.$i.out 2> vp-dc.r200.$i.err
done

Where nc.r200.fa.$i is a fasta file with 1 of the 200 sequences from
nc.r200.fa, and nc.r200.$i.csv has the metadata for that
sequence. These files are included in the tarball in the
seq-files/vapid-indi-files directory. 

VAPiD was run a single sequence at a time like this because if no
reference sequence is found for the current input sequence it is
processing, the program will exit (in my hands). So to make it run on
all 200 sequences, each sequence was separated to its own file.

VAPiD was not run on the partial sequence sets (np and dp) because it
was designed for complete genomes as discussed in the VADR paper.

VAPiD was not run on the full sets of test sequences. 

Results were processed as explained in
README-processing-test-results.txt

============================
INSTALLING AND RUNNING VIGOR
============================

Downloaded VIGOR3.r20.tgz from:

https://sourceforge.net/projects/jcvi-vigor/ 

Which I reached from:
https://www.jcvi.org/gcid/project/vigor -> 'Vigor Repository' (under resources)

Unpacked gzipped tarball
$ tar xfz VIGOR3.r20.tgz 

And followed instructions in the file VIGOR3/README.txt

To get VIGOR to run in my hands I had to make the following change to
line 55 of VIGOR3/prod3/db.pm from:

my $dbh = DBI->connect( $dsn, "", "", {PrintError=>1,RaiseError=>1, sqlite_open_flags => DBD::SQLite::OPEN_READONLY} );

to

my $dbh = DBI->connect( $dsn, "", "", {PrintError=>1,RaiseError=>1, sqlite_open_flags => "DBD::SQLite::OPEN_READONLY"} );

(added quotes around DBD::SQLite::OPEN_READONLY)

--

VIGOR was run on the Norovirus 200 sequence sets on a single processor
with the following commands:

$ VIGOR3.pl -i np.r200.fa -o vg-np.r200/vg-np.r200 > vg-np.r200.out
$ VIGOR3.pl -i nc.r200.fa -o vg-nc.r200/vg-nc.r200 > vg-nc.r200.out

VIGOR was not run on the Dengue virus sequences because it does not
contain a Dengue reference library as discussed in the VADR paper. 

VIGOR was not run on the full sets of test sequences. 

Results were processed as explained in
README-processing-test-results.txt

----------------------------------------
Questions, contact
----------------------------------------
