## Supplemental material tarball for "VADR: validation and annotation of virus sequence submissions to GenBank": acclist2mp-and-cds.pl

```
my $usage = "perl acclist2mp-and-cds.pl <list of accessions> <'mp' or 'cds'>";
if(scalar(@ARGV) != 2) { die $usage; }

my ($list_file, $mp_or_cds) = (@ARGV);
my $type = "CDS";
if($mp_or_cds eq "mp") { 
  $type = "mat_peptide";
}
elsif($mp_or_cds ne "cds") { 
  die $usage;
}

my $n = 0;
my $chunk = 30;
open(IN, $list_file) || die "ERROR unable to open $list_file for reading";
while($line = <IN>) { 
  chomp $line;
  if($n > 0) { 
    $query .= "," . $line;
  }
  else { 
    $query = $line;
  }
  $n++;
  if($n == $chunk) { 
    my $cmd = "esearch -db nuccore -query \"$query\" | efetch -format gpc | xtract -insd $type INSDFeature_location";
    #printf("running $cmd\n");
    system($cmd);
    $n = 0;
    $query = "";
  }
}
close(IN);
if($n > 0) { 
  my $cmd = "esearch -db nuccore -query \"$query\" | efetch -format gpc | xtract -insd $type INSDFeature_location";
  #printf("running $cmd\n");
  system($cmd);
}
```
