## Supplemental material tarball for "VADR: validation and annotation of virus sequence submissions to GenBank": combine-seqsums.pl

```
#!/usr/bin/env perl
use strict;

my $usage = "perl combine-seqsums.pl <.seqsum file 1> <name for file 1, e.g. \"vadr\"> <.seqsum file 2> <name for file 2, e.g. \"vigor\">\n";
$usage   .= " OR\n";
$usage   .= "perl combine-seqsums.pl <.seqsum file 1> <name for file 1, e.g. \"vadr\"> <.seqsum file 2> <name for file 2, e.g. \"vigor\"> <.seqsum file 3> <name for file 3, e.g. \"vapid\"\n";
$usage   .= "\nNOTE: if one file is for vigor, it must be the second file\n";

my $seqsum1_file = undef;
my $prefix1      = undef;
my $lc_prefix1   = undef;

my $seqsum2_file = undef;
my $prefix2      = undef;
my $lc_prefix2   = undef;

my $seqsum3_file = undef;
my $prefix3      = undef;
my $lc_prefix3   = undef;

my $do_2 = 0;
my $do_3 = 0;
if(scalar(@ARGV) == 4) { 
  $do_2 = 1;  
  ($seqsum1_file, $prefix1, $seqsum2_file, $prefix2) = (@ARGV);
}
elsif(scalar(@ARGV) == 6) { 
  $do_3 = 1; 
  ($seqsum1_file, $prefix1, $seqsum2_file, $prefix2, $seqsum3_file, $prefix3) = (@ARGV);
}
else {
  die $usage; 
}

# vigor has to be file 2
$lc_prefix1 = $prefix1;
$lc_prefix1 =~ tr/A-Z/a-z/;
if($lc_prefix1 eq "vigor") { die "ERROR vigor file must be file 2\n"; }

$lc_prefix2 = $prefix2;
$lc_prefix2 =~ tr/A-Z/a-z/;

if($do_3) { 
  $lc_prefix3 = $prefix3;
  $lc_prefix3 =~ tr/A-Z/a-z/;
  if($lc_prefix3 eq "vigor") { die "ERROR vigor file must be file 2\n"; }
}

my $seqname;
my $line; # a line of a file
my %pf1_H = (); # key is sequence name, value is PASS or FAIL
my %pf2_H = (); # key is sequence name, value is PASS or FAIL
my %pf3_H = (); # key is sequence name, value is PASS or FAIL
my %pfreason1_H = (); # key is sequence name, value is reason for FAILure or "NULL" if it passes
my %pfreason2_H = (); # key is sequence name, value is reason for FAILure or "NULL" if it passes
my %pfreason3_H = (); # key is sequence name, value is reason for FAILure or "NULL" if it passes
my @seqname_A = (); # array of sequence names in order they were read in
my %seqname_ct_H = (); # number of times seqname was read 

open(SEQSUM1, $seqsum1_file) || die "ERROR unable to open $seqsum1_file for reading";
while($line = <SEQSUM1>) { 
  chomp $line;
  my @el_A = split(/\s+/, $line);
  if($el_A[0] =~ /^seq\:(\S+)/) { 
    $seqname = $1;
    push(@seqname_A, $seqname);
    if(defined $seqname_ct_H{$seqname}) { die "ERROR read sequence $seqname twice in file 1"; }
    $seqname_ct_H{$seqname} = 1;
  }
  else { die "ERROR first token not seq seqsum1 file line:$line\n"; }

  if($el_A[1] =~ /^pf\:(\S+)/) { 
    my $pf = $1;
    $pf1_H{$seqname} = "pf-" . $lc_prefix1 . ":" . $1;
  }
  else { die "ERROR 2nd token not pf seqsum1 file line:$line\n"; }

  if($el_A[2] =~ /^pfreason\:(\S+)/) { 
    my $pfreason = $1;
    $pfreason1_H{$seqname} = "pfreason-" . $lc_prefix1 . ":" . $1;
  }
  else { die "ERROR 3rd token not pfreason seqsum1 file line:$line\n"; }
}
close(SEQSUM1);

open(SEQSUM2, $seqsum2_file) || die "ERROR unable to open $seqsum2_file for reading";
while($line = <SEQSUM2>) { 
  chomp $line;
  my @el_A = split(/\s+/, $line);
  if($el_A[0] =~ /^seq\:(\S+)/) { 
    $seqname = $1;
    if(! defined $seqname_ct_H{$seqname}) { die "ERROR read sequence $seqname from file 2 but not file 1"; }
    if($seqname_ct_H{$seqname} != 1) { die "ERROR read sequence $seqname more than once in file 2"; }
    $seqname_ct_H{$seqname} = 2;
  }
  else { die "ERROR first token not seq seqsum2 file line:$line\n"; }

  if($el_A[1] =~ /^pf\:(\S+)/) { 
    my $pf = $1;
    $pf2_H{$seqname} = "pf-" . $lc_prefix2 . ":" . $1;
  }
  else { die "ERROR 2nd token not pf seqsum2 file line:$line\n"; }

  if($el_A[2] =~ /^pfreason\:(\S+)/) { 
    my $pfreason = $1;
    $pfreason2_H{$seqname} = "pfreason-" . $lc_prefix2 . ":" . $1;
  }
  else { die "ERROR 3rd token not pfreason seqsum2 file line:$line\n"; }
}
close(SEQSUM2);

# deal with fact that vigor may not have all seqs in its seqsum file
if($lc_prefix2) { 
  # vigor may not have all files in the .seqsum file (derived from the .at file)
  foreach $seqname (@seqname_A) { 
    if(! exists $pf2_H{$seqname}) { 
      # this seq was in file 1 but not file 2
      $pf2_H{$seqname}       = "pf-" . $lc_prefix2 . ":" . "FAIL";
      $pfreason2_H{$seqname} = "pfreason-" . $lc_prefix2 . ":" . "NOANNOTATION"; 
      $seqname_ct_H{$seqname} = 2;
    }
  }
}

if($do_3) { 
  open(SEQSUM3, $seqsum3_file) || die "ERROR unable to open $seqsum3_file for reading";
  while($line = <SEQSUM3>) { 
    chomp $line;
    my @el_A = split(/\s+/, $line);
    if($el_A[0] =~ /^seq\:(\S+)/) { 
      $seqname = $1;
      if(! defined $seqname_ct_H{$seqname}) { die "ERROR read sequence $seqname from file 3 but not file 1"; }
      if($seqname_ct_H{$seqname} != 2) { die "ERROR read sequence $seqname in file 3, but it's not the 3rd time I've seen this seqname"; }
      $seqname_ct_H{$seqname} = 3;
    }
    else { die "ERROR first token not seq seqsum3 file line:$line\n"; }
    
    if($el_A[1] =~ /^pf\:(\S+)/) { 
      my $pf = $1;
      $pf3_H{$seqname} = "pf-" . $lc_prefix3 . ":" . $1;
    }
    else { die "ERROR 2nd token not pf seqsum2 file line:$line\n"; }
    
    if($el_A[2] =~ /^pfreason\:(\S+)/) { 
      my $pfreason = $1;
      $pfreason3_H{$seqname} = "pfreason-" . $lc_prefix3 . ":" . $1;
    }
    else { die "ERROR 3rd token not pfreason seqsum3 file line:$line\n"; }
  }
  close(SEQSUM3);
}    

foreach $seqname (@seqname_A) { 
  if($do_2) { 
    if($seqname_ct_H{$seqname} != 2) { die "ERROR didn't read $seqname in both files"; }
    printf ("seq:%-10s %s %s %s %s\n", $seqname, $pf1_H{$seqname}, $pfreason1_H{$seqname}, $pf2_H{$seqname}, $pfreason2_H{$seqname}); 
  }
  elsif($do_3) { 
    if($seqname_ct_H{$seqname} != 3) { die "ERROR didn't read $seqname in all 3 files"; }
    printf ("seq:%-10s %s %s %s %s %s %s\n", $seqname, $pf1_H{$seqname}, $pfreason1_H{$seqname}, $pf2_H{$seqname}, $pfreason2_H{$seqname}, $pf3_H{$seqname}, $pfreason3_H{$seqname}); 
  }
}
```
