## Supplemental material tarball for "VADR: validation and annotation of virus sequence submissions to GenBank": combine-summaries-for-tbl.sh

```
#!/bin/bash
# table 1: pass/fail for each program on each dataset
for a in nc; do 
    echo -n "$a vadr  pass "
    grep pf-vadr:PASS $a.vadr.vigor.vapid.combseqsum | wc -l
    echo -n "$a vadr  fail "
    grep pf-vadr:FAIL $a.vadr.vigor.vapid.combseqsum | wc -l
    echo -n "$a vapid pass "
    grep pf-vapid:PASS $a.vadr.vigor.vapid.combseqsum | wc -l
    echo -n "$a vapid fail "
    grep pf-vapid:FAIL $a.vadr.vigor.vapid.combseqsum | wc -l
    echo -n "$a vigor pass "
    grep pf-vigor:PASS $a.vadr.vigor.vapid.combseqsum | wc -l
    echo -n "$a vigor fail "
    grep pf-vigor:FAIL $a.vadr.vigor.vapid.combseqsum | wc -l
    echo ""
done
for a in dc; do 
    echo -n "$a vadr  pass "
    grep pf-vadr:PASS $a.vadr.vapid.combseqsum | wc -l
    echo -n "$a vadr  fail "
    grep pf-vadr:FAIL $a.vadr.vapid.combseqsum | wc -l
    echo -n "$a vapid pass "
    grep pf-vapid:PASS $a.vadr.vapid.combseqsum | wc -l
    echo -n "$a vapid fail "
    grep pf-vapid:FAIL $a.vadr.vapid.combseqsum | wc -l
    echo "$a vigor pass -"
    echo "$a vigor fail -"
    echo ""
done
for a in np; do 
    echo -n "$a vadr  pass "
    grep pf-vadr:PASS $a.vadr.vigor.combseqsum | wc -l
    echo -n "$a vadr  fail "
    grep pf-vadr:FAIL $a.vadr.vigor.combseqsum | wc -l
    echo "$a vapid pass -"
    echo "$a vapid fail -"
    echo -n "$a vigor pass "
    grep pf-vigor:PASS $a.vadr.vigor.combseqsum | wc -l
    echo -n "$a vigor fail "
    grep pf-vigor:FAIL $a.vadr.vigor.combseqsum | wc -l
    echo ""
done
for a in dp; do 
    echo -n "$a vadr  pass "
    grep pf:PASS vadr.$a.seqsum | wc -l
    echo -n "$a vadr  fail "
    grep pf:FAIL vadr.$a.seqsum | wc -l
    echo "$a vapid pass -"
    echo "$a vapid fail -"
    echo "$a vigor pass -"
    echo "$a vigor fail -"
    echo ""
done

    echo ""
# table 2: vadr vs vapid and vadr vs vigor
for a in nc; do 
    echo -n "$a vadr-vapid   pass-pass "
    grep pf-vadr:PASS $a.vadr.vigor.vapid.combseqsum | grep pf-vapid:PASS | wc -l
    echo -n "$a vadr-vapid   fail-fail "
    grep pf-vadr:FAIL $a.vadr.vigor.vapid.combseqsum | grep pf-vapid:FAIL | wc -l
    echo -n "$a vadr-vapid   pass-fail "
    grep pf-vadr:PASS $a.vadr.vigor.vapid.combseqsum | grep pf-vapid:FAIL | wc -l
    echo -n "$a vadr-vapid   fail-pass "
    grep pf-vadr:FAIL $a.vadr.vigor.vapid.combseqsum | grep pf-vapid:PASS | wc -l
    echo ""
done
for a in dc; do 
    echo -n "$a vadr-vapid   pass-pass "
    grep pf-vadr:PASS $a.vadr.vapid.combseqsum | grep pf-vapid:PASS | wc -l
    echo -n "$a vadr-vapid   fail-fail "
    grep pf-vadr:FAIL $a.vadr.vapid.combseqsum | grep pf-vapid:FAIL | wc -l
    echo -n "$a vadr-vapid   pass-fail "
    grep pf-vadr:PASS $a.vadr.vapid.combseqsum | grep pf-vapid:FAIL | wc -l
    echo -n "$a vadr-vapid   fail-pass "
    grep pf-vadr:FAIL $a.vadr.vapid.combseqsum | grep pf-vapid:PASS | wc -l
    echo ""
done

for a in nc; do 
    echo -n "$a vadr-vigor   pass-pass "
    grep pf-vadr:PASS $a.vadr.vigor.vapid.combseqsum | grep pf-vigor:PASS | wc -l
    echo -n "$a vadr-vigor   fail-fail "
    grep pf-vadr:FAIL $a.vadr.vigor.vapid.combseqsum | grep pf-vigor:FAIL | wc -l
    echo -n "$a vadr-vigor   pass-fail "
    grep pf-vadr:PASS $a.vadr.vigor.vapid.combseqsum | grep pf-vigor:FAIL | wc -l
    echo -n "$a vadr-vigor   fail-pass "
    grep pf-vadr:FAIL $a.vadr.vigor.vapid.combseqsum | grep pf-vigor:PASS | wc -l
    echo ""
done
for a in np; do 
    echo -n "$a vadr-vigor   pass-pass "
    grep pf-vadr:PASS $a.vadr.vigor.combseqsum | grep pf-vigor:PASS | wc -l
    echo -n "$a vadr-vigor   fail-fail "
    grep pf-vadr:FAIL $a.vadr.vigor.combseqsum | grep pf-vigor:FAIL | wc -l
    echo -n "$a vadr-vigor   pass-fail "
    grep pf-vadr:PASS $a.vadr.vigor.combseqsum | grep pf-vigor:FAIL | wc -l
    echo -n "$a vadr-vigor   fail-pass "
    grep pf-vadr:FAIL $a.vadr.vigor.combseqsum | grep pf-vigor:PASS | wc -l
    echo ""
done
```
