## Supplemental material tarball for "VADR: validation and annotation of virus sequence submissions to GenBank": combine-summaries.sh

```
#!/bin/bash
# nc: compare all 3
for a in nc; do 
    echo "------------------"
    echo "Norovirus-complete"
    echo "------------------"
    echo ""
    for b in vadr vigor vapid; do 
        echo -n "num PASS $b " 
        grep pf:PASS $b.$a.seqsum | wc -l
    done
    echo ""

    perl $VADRPAPER_SCRIPTSDIR/combine-seqsums.pl vadr.$a.seqsum vadr vigor.$a.seqsum vigor vapid.$a.seqsum vapid > $a.vadr.vigor.vapid.combseqsum
    # 3/3 pass
    # num vadr-pass vigor-pass vapid-pass
    grep pf-vadr:PASS $a.vadr.vigor.vapid.combseqsum | grep pf-vigor:PASS | grep pf-vapid:PASS > $a.vadr-pass.vigor-pass.vapid-pass.list
    wc -l $a.vadr-pass.vigor-pass.vapid-pass.list

    # 2/3 pass
    # num vadr-pass vigor-pass vapid-fail
    grep pf-vadr:PASS $a.vadr.vigor.vapid.combseqsum | grep pf-vigor:PASS | grep pf-vapid:FAIL > $a.vadr-pass.vigor-pass.vapid-fail.list
    wc -l $a.vadr-pass.vigor-pass.vapid-fail.list
    # num vadr-pass vigor-fail vapid-pass
    grep pf-vadr:PASS $a.vadr.vigor.vapid.combseqsum | grep pf-vigor:FAIL | grep pf-vapid:PASS > $a.vadr-pass.vigor-fail.vapid-pass.list
    wc -l $a.vadr-pass.vigor-fail.vapid-pass.list
    # num vadr-fail vigor-pass vapid-pass
    grep pf-vadr:FAIL $a.vadr.vigor.vapid.combseqsum | grep pf-vigor:PASS | grep pf-vapid:PASS > $a.vadr-fail.vigor-pass.vapid-pass.list
    wc -l $a.vadr-fail.vigor-pass.vapid-pass.list

    # 1/3 pass
    # num vadr-pass vigor-fail vapid-fail
    grep pf-vadr:PASS $a.vadr.vigor.vapid.combseqsum | grep pf-vigor:FAIL | grep pf-vapid:FAIL > $a.vadr-pass.vigor-fail.vapid-fail.list
    wc -l $a.vadr-pass.vigor-fail.vapid-fail.list
    # num vadr-fail vigor-pass vapid-fail
    grep pf-vadr:FAIL $a.vadr.vigor.vapid.combseqsum | grep pf-vigor:PASS | grep pf-vapid:FAIL > $a.vadr-fail.vigor-pass.vapid-fail.list
    wc -l $a.vadr-fail.vigor-pass.vapid-fail.list
    # num vadr-fail vigor-fail vapid-pass
    grep pf-vadr:FAIL $a.vadr.vigor.vapid.combseqsum | grep pf-vigor:FAIL | grep pf-vapid:PASS > $a.vadr-fail.vigor-fail.vapid-pass.list
    wc -l $a.vadr-fail.vigor-fail.vapid-pass.list

    # 0/3 pass
    # num vadr-fail vigor-fail vapid-faail
    grep pf-vadr:FAIL $a.vadr.vigor.vapid.combseqsum | grep pf-vigor:FAIL | grep pf-vapid:FAIL > $a.vadr-fail.vigor-fail.vapid-fail.list
    wc -l $a.vadr-fail.vigor-fail.vapid-fail.list

done

echo ""
# np: compare vadr and vigor
for a in np; do 
    echo "-----------------"
    echo "Norovirus-partial"
    echo "-----------------"
    echo ""
    for b in vadr vigor; do 
        echo -n "num PASS $b " 
        grep pf:PASS $b.$a.seqsum | wc -l
    done
    echo ""

    perl $VADRPAPER_SCRIPTSDIR/combine-seqsums.pl vadr.$a.seqsum vadr vigor.$a.seqsum vigor > $a.vadr.vigor.combseqsum

    # 2/2 pass
    # num vadr-pass vigor-pass
    grep pf-vadr:PASS $a.vadr.vigor.combseqsum | grep pf-vigor:PASS > $a.vadr-pass.vigor-pass.list
    wc -l $a.vadr-pass.vigor-pass.list

    # 1/2 pass
    # num vadr-pass vigor-fail
    grep pf-vadr:PASS $a.vadr.vigor.combseqsum | grep pf-vigor:FAIL > $a.vadr-pass.vigor-fail.list
    wc -l $a.vadr-pass.vigor-fail.list
    # num vadr-fail vigor-pass
    grep pf-vadr:FAIL $a.vadr.vigor.combseqsum | grep pf-vigor:PASS > $a.vadr-fail.vigor-pass.list
    wc -l $a.vadr-fail.vigor-pass.list

    # 0/2 pass
    # num vadr-fail vigor-fail
    grep pf-vadr:FAIL $a.vadr.vigor.combseqsum | grep pf-vigor:FAIL > $a.vadr-fail.vigor-fail.list
    wc -l $a.vadr-fail.vigor-fail.list
done

echo ""
# dc: compare vadr and vapid
for a in dc; do 
    echo "---------------"
    echo "Dengue-complete"
    echo "---------------"
    echo ""
    for b in vadr vapid; do 
        echo -n "num PASS $b " 
        grep pf:PASS $b.$a.seqsum | wc -l
    done
    echo ""

    perl $VADRPAPER_SCRIPTSDIR/combine-seqsums.pl vadr.$a.seqsum vadr vapid.$a.seqsum vapid > $a.vadr.vapid.combseqsum

    # 2/2 pass
    # num vadr-pass vapid-pass
    grep pf-vadr:PASS $a.vadr.vapid.combseqsum | grep pf-vapid:PASS > $a.vadr-pass.vapid-pass.list
    wc -l $a.vadr-pass.vapid-pass.list

    # 1/2 pass
    # num vadr-pass vapid-fail
    grep pf-vadr:PASS $a.vadr.vapid.combseqsum | grep pf-vapid:FAIL > $a.vadr-pass.vapid-fail.list
    wc -l $a.vadr-pass.vapid-fail.list
    # num vadr-fail vapid-pass
    grep pf-vadr:FAIL $a.vadr.vapid.combseqsum | grep pf-vapid:PASS > $a.vadr-fail.vapid-pass.list
    wc -l $a.vadr-fail.vapid-pass.list

    # 0/2 pass
    # num vadr-fail vapid-fail
    grep pf-vadr:FAIL $a.vadr.vapid.combseqsum | grep pf-vapid:FAIL > $a.vadr-fail.vapid-fail.list
    wc -l $a.vadr-fail.vapid-fail.list
done
```
