## Supplemental material tarball for "VADR: validation and annotation of virus sequence submissions to GenBank": compare-two-ftrsums.pl

```
#!/usr/bin/env perl
use strict;

my $usage = "perl compare-two-ftrsums.pl <.ftrsum file 1> <.ftrsum file 2> <'all' or 'pass' (for only counting features in seqs that pass both methods)>\n";

if(scalar(@ARGV) != 3) { 
  die $usage;
}
my ($ftrsum1_file, $ftrsum2_file, $all_or_pass) = (@ARGV);

my $do_all = undef;
if($all_or_pass eq "all") { 
  $do_all = 1;
}
elsif($all_or_pass eq "pass") { 
  $do_all = 0;
}
if(! defined $do_all) { 
  die $usage;
}

my $line;
my $seq;
my $product;
my %coords1_H = ();
my %coords2_H = ();
my %ref1_H = ();
my %ref2_H = ();
my %pf1_H = ();
my %pf2_H = ();

open(FTRSUM1, $ftrsum1_file) || die "ERROR unable to open $ftrsum1_file for reading";
while($line = <FTRSUM1>) { 
  chomp $line;
  my @el_A = split(/\s+/, $line);
  foreach my $el (@el_A) { 
    if($el =~ /^coords\:(\S+)$/) { 
      $coords1_H{$seq.":".$product} = $1;
    }
    elsif($el =~ /^([^\:]+)\:([^\:]+)$/) { 
      my ($key, $value) = ($1, $2); 
      if($key eq "seq") { 
        $seq = $value; 
      }
      elsif($key eq "product") { 
        $product = $value; 
#        $product =~ s/^genome\_//;
        $product =~ s/^capsid\_protein\_//;
      }
      elsif($key eq "ref") { 
        $ref1_H{$seq.":".$product} = $value; 
      }
      elsif($key eq "pf") { 
        $pf1_H{$seq.":".$product} = $value; 
      }
    }
    else { die "ERROR unable to parse element $el in line:\n$line"; }
  }
}
close(FTRSUM1);

open(FTRSUM2, $ftrsum2_file) || die "ERROR unable to open $ftrsum2_file for reading";
while($line = <FTRSUM2>) { 
  chomp $line;
  my @el_A = split(/\s+/, $line);
  foreach my $el (@el_A) { 
    if($el =~ /^coords\:(\S+)$/) { 
      $coords2_H{$seq.":".$product} = $1;
    }
    elsif($el =~ /^([^\:]+)\:([^\:]+)$/) { 
      my ($key, $value) = ($1, $2); 
      if($key eq "seq") { 
        $seq = $value; 
      }
      elsif($key eq "product") { 
        $product = $value; 
#        $product =~ s/^genome\_//;
        $product =~ s/^capsid\_protein\_//;
      }
      elsif($key eq "ref") { 
        $ref2_H{$seq.":".$product} = $value; 
      }
      elsif($key eq "pf") { 
        $pf2_H{$seq.":".$product} = $value; 
      }
    }
    else { die "ERROR unable to parse element $el in line:\n$line"; }
  }
}
close(FTRSUM2);

foreach my $seqproduct (sort keys (%ref1_H)) { 
  if(($do_all) || 
     ((defined $pf1_H{$seqproduct}) && (defined $pf2_H{$seqproduct}) && 
      ($pf1_H{$seqproduct} eq "PASS") && $pf2_H{$seqproduct} eq "PASS")) {  
    if(defined $ref2_H{$seqproduct}) { 
      my $coords1 = $coords1_H{$seqproduct};
      my $coords2 = $coords2_H{$seqproduct};
      $coords1 =~ s/\>//;
      $coords1 =~ s/\<//;
      $coords2 =~ s/\>//;
      $coords2 =~ s/\<//;
      if($ref1_H{$seqproduct} eq $ref2_H{$seqproduct}) { 
        if($coords1 eq $coords2) { 
          printf("PRODUCT-MATCH-REF-MATCH-COORDS-MATCH $seqproduct ref:$ref1_H{$seqproduct} coords: $coords1_H{$seqproduct} $coords2_H{$seqproduct}\n");
        }
        else { 
          printf("PRODUCT-MATCH-REF-MATCH-COORDS-MISMATCH $seqproduct ref:$ref1_H{$seqproduct} coords: $coords1_H{$seqproduct} $coords2_H{$seqproduct}\n");
        }
      }
      else { 
        if($coords1 eq $coords2) { 
          printf("PRODUCT-MATCH-REF-MISMATCH-COORDS-MATCH $seqproduct ref:$ref1_H{$seqproduct} coords: $coords1_H{$seqproduct} $coords2_H{$seqproduct}\n");
        }
        else { 
          printf("PRODUCT-MATCH-REF-MISMATCH-COORDS-MISMATCH $seqproduct ref:$ref1_H{$seqproduct} coords: $coords1_H{$seqproduct} $coords2_H{$seqproduct}\n");
        }
      }
    }
    else { 
      printf("PRODUCT-MISMATCH $seqproduct\n");
    }
  }
}
```
