## Supplemental material tarball for "VADR: validation and annotation of virus sequence submissions to GenBank": count-ambigs.pl

```
my $seqname = undef;
my $seq = "";
my $cur_nambig = 0;
my $new_seqname = "";
while($line = <>) { 
  chomp $line;
  if($line =~ /^\>(\S+)/) { 
    $new_seqname = $1;
    if($seq ne "") { 
      $cur_nambig = 0;
      $seq =~ tr/a-z/A-Z/;
      $seq =~ s/U/T/;
      my @seq_A = split("", $seq);
      my $cur_len = scalar(@seq_A);
      for(my $i = 0; $i < $cur_len; $i++) { 
        if(($seq_A[$i] ne "A") && 
           ($seq_A[$i] ne "C") && 
           ($seq_A[$i] ne "G") && 
           ($seq_A[$i] ne "T")) { 
          $cur_nambig++;
        }
      }
      printf("%s %5d %5d %.4f\n", $seqname, $cur_nambig, $cur_len, ($cur_nambig/$cur_len)); 
    }
    $seqname = $new_seqname;
    $seq = "";
  }
  else { 
    $seq .= $line;
  }
}
if($seq ne "") { 
  if($seq ne "") { 
    $cur_nambig = 0;
    $seq =~ tr/a-z/A-Z/;
    $seq =~ s/U/T/;
    my @seq_A = split("", $seq);
    my $cur_len = scalar(@seq_A);
    for(my $i = 0; $i < $cur_len; $i++) { 
      if(($seq_A[$i] ne "A") && 
         ($seq_A[$i] ne "C") && 
         ($seq_A[$i] ne "G") && 
         ($seq_A[$i] ne "T")) { 
        $cur_nambig++;
        $cur_nambig++;
      }
    }
    printf("%s %5d %5d %.4f\n", $seqname, $cur_nambig, $cur_len, ($cur_nambig/$cur_len)); 
  }
  $seqname = $new_seqname;
  $seq = "";
}
```
