## Supplemental material tarball for "VADR: validation and annotation of virus sequence submissions to GenBank": genbank-count-cds-and-mp.sh

```
#!/bin/bash
for a in np dp nc dc; do
    echo $a 
    echo -n "total number seqs: "
    wc -l $VADRPAPER_SUPPDIR/seq-files/$a.list
    echo -n "num seqs with >= 1 CDS annotated: "
    cat $a.cds.out | awk '{ print $1 }' | sort | uniq | wc -l
    echo -n "num seqs with >= 1 mat_peptide annotated: "
    cat $a.mp.out | awk '{ print $1 }' | sort | uniq | wc -l
    echo ""
done
```
