## Supplemental material tarball for "VADR: validation and annotation of virus sequence submissions to GenBank": genbank-fetch-cds-and-mp.sh

```
#!/bin/bash
for a in np dp nc dc; do
    perl $VADRPAPER_SCRIPTSDIR/acclist2mp-and-cds.pl $VADRPAPER_SUPPDIR/seq-files/$a.list cds > $a.cds.out
    perl $VADRPAPER_SCRIPTSDIR/acclist2mp-and-cds.pl $VADRPAPER_SUPPDIR/seq-files/$a.list mp > $a.mp.out
done
```
