## Supplemental material tarball for "VADR: validation and annotation of virus sequence submissions to GenBank": longest-N-stretch.pl

```
my $seqname = undef;
my $seq = "";
my $cur_n = 0;
my $max_n = 0;
my $new_seqname = "";
while($line = <>) { 
  chomp $line;
  if($line =~ /^\>(\S+)/) { 
    $new_seqname = $1;
    if($seq ne "") { 
      $cur_n = 0;
      $max_n = 0;
      $seq =~ tr/a-z/A-Z/;
      my @seq_A = split("", $seq);
      my $n = scalar(@seq_A);
      for(my $i = 0; $i < $n; $i++) { 
        if($seq_A[$i] eq "N") { 
          $cur_n++;
          if($cur_n > $max_n) { $max_n = $cur_n; }
        }
        else { 
          $cur_n = 0;
        }
      }
      printf("%s %5d\n", $seqname, $max_n); 
    }
    $seqname = $new_seqname;
    $seq = "";
  }
  else { 
    $seq .= $line;
  }
}
if($seq ne "") { 
  $cur_n = 0;
  $max_n = 0;
  $seq =~ tr/a-z/A-Z/;
  my @seq_A = split("", $seq);
  my $n = scalar(@seq_A);
  for(my $i = 0; $i < $n; $i++) { 
    if($seq_A[$i] eq "N") { 
      $cur_n++;
      if($cur_n > $max_n) { $max_n = $cur_n; }
    }
    else { 
      $cur_n = 0;
    }
  }
  printf("%s %5d\n", $seqname, $max_n); 
}  
```
