## Supplemental material tarball for "VADR: validation and annotation of virus sequence submissions to GenBank": make-summaries.sh

```
#!/bin/bash
#vapid
for a in nc dc; do 
    if [ -e vapid.$a.seqsum ] 
    then 
        rm vapid.$a.seqsum
    fi
    for ((i = 1; i <= 200; i++)); do
        perl $VADRPAPER_SCRIPTSDIR/vapid-out-and-errorsummary-to-seqsum.pl $VADRPAPER_VAPIDDIR/vapid-r200indi-out/vp-$a.r200.$i/vp-$a.r200.$i.out $VADRPAPER_VAPIDDIR/vapid-r200indi-out/vp-$a.r200.$i $VADRPAPER_SUPPDIR/seq-files/vapid-indi-files/$a.r200.$i.namelist >> vapid.$a.seqsum 
    done
done

#vigor
for a in nc np; do 
    perl $VADRPAPER_SCRIPTSDIR/vigor-at-to-seqsum-and-ftrsum.pl $VADRPAPER_VIGORDIR/vigor-r200-out/vg-$a.r200/vg-$a.r200.at vigor.$a
done

#vadr
for a in np nc; do 
    perl $VADRPAPER_SCRIPTSDIR/vadr-sqa-and-ftr-to-seqsum-and-ftrsum.pl $VADRPAPER_VADRDIR/vadr-r200-out/va-$a.r200/va-$a.r200.vadr.sqa $VADRPAPER_VADRDIR/vadr-r200-out/va-$a.r200/va-$a.r200.vadr.ftr $VADRPAPER_SUPPDIR/vadr.noro.map vadr.$a
done
for a in dp dc; do 
    perl $VADRPAPER_SCRIPTSDIR/vadr-sqa-and-ftr-to-seqsum-and-ftrsum.pl $VADRPAPER_VADRDIR/vadr-r200-out/va-$a.r200/va-$a.r200.vadr.sqa $VADRPAPER_VADRDIR/vadr-r200-out/va-$a.r200/va-$a.r200.vadr.ftr $VADRPAPER_SUPPDIR/vadr.dengue.map vadr.$a
done
```
