## Supplemental material tarball for "VADR: validation and annotation of virus sequence submissions to GenBank": set-environment.sh

```
#!/bin/bash

# set VADRPAPER_SUPPDIR to the path you unpacked the supplementary data tarball 
export VADRPAPER_SUPPDIR="$INF/notebook/19_1003_vadr_mscript_tests3/m-vadr/vadr-1.0-paper-supplementary-material"

# set VADRPAPER_VADRDIR to the path where you ran vadr as explained here in the file 
# 00NOTES-running-tests.txt included in the supplementary data tarball
export VADRPAPER_VADRDIR="$INF/notebook/19_1003_vadr_mscript_tests3/vadr-dir"

# set VADRPAPER_VAPIDDIR to the path where you ran vapid as explained here in the file 
# 00NOTES-running-tests.txt included in the supplementary data tarball
export VADRPAPER_VAPIDDIR="$INF/notebook/19_1003_vadr_mscript_tests3/vapid-dir"

# set VADRPAPER_VIGORDIR to the path where you ran vigor as explained here in the file 
# 00NOTES-running-tests.txt included in the supplementary data tarball
export VADRPAPER_VIGORDIR="$INF/notebook/19_1003_vadr_mscript_tests3/vigor-dir"

# do not edit the next line
export VADRPAPER_SCRIPTSDIR="$VADRPAPER_SUPPDIR/scripts"
```
