## Supplemental material tarball for "VADR: validation and annotation of virus sequence submissions to GenBank": vadr-count-rnas-full.sh

```
#!/bin/bash
DIR="$VADRPAPER_VADRDIR/vadr-full-out"
for a in dp dc; do 
    echo ""
    echo -n "$a total num passing seqs: "
    grep -v ^\# $DIR/va-$a/va-$a.vadr.sqa | grep PASS | wc -l
    for b in ncRNA stem_loop; do 
        echo -n "$a $b total num features: "
        grep PASS $DIR/va-$a/va-$a.vadr.ftr | grep $b | wc -l
        echo -n "$a $b num seqs with >= 1: "
        grep PASS $DIR/va-$a/va-$a.vadr.ftr | grep $b | awk '{ print $2 }' | sort | uniq -c | wc -l
        echo -n "$a $b seq with max num: "
        grep PASS $DIR/va-$a/va-$a.vadr.ftr | grep $b | awk '{ print $2 }' | sort | uniq -c | sort -rnk 1 | head -n1
        echo -n "$a $b seq with min num: "
        grep PASS $DIR/va-$a/va-$a.vadr.ftr | grep $b | awk '{ print $2 }' | sort | uniq -c | sort -rnk 1 | tail -n1
    done
done

        
```
