## Supplemental material tarball for "VADR: validation and annotation of virus sequence submissions to GenBank": vadr-make-alertcounts-table.pl

```
#!/usr/bin/env perl
use strict;

my $usage = "perl vadr-make-alertcounts-table.pl <v-annotate.pl --alt_list output> <'full' or 'test'>";

if(scalar(@ARGV) != 2) { 
  die $usage;
}
my ($alt_list, $full_or_test) = (@ARGV);


if(! exists($ENV{"VADRPAPER_VADRDIR"})) { 
  die "ERROR, the environment variable VADRPAPER_VADRDIR is not set";
}
my $results_dir = $ENV{"VADRPAPER_VADRDIR"};

my @sets_A = ("nc", "np", "dc", "dp");
my $do_full = 0;
if($full_or_test eq "full") { 
#  $results_dir = "../../19_0705_vadr_mscript_tests/vadr-dir/vadr-out-allseqs-longnames-20190709-1";
  $results_dir .= "/vadr-full-out/";
  $do_full = 1;
}
elsif($full_or_test eq "test") { 
  $results_dir .= "/vadr-r200-out/";
}
if(! -d $results_dir) { die "ERROR results dir $results_dir does not exist"; }

my $code;
my $desc;
my @code_A = ();
my %desc_H = ();

# hard-coded non-fatal alerts
my %nonfatal_H = ();
$nonfatal_H{"qstsbgrp"} = 1;
$nonfatal_H{"qstgroup"} = 1;
$nonfatal_H{"indfclas"} = 1;
$nonfatal_H{"lowscore"} = 1;
$nonfatal_H{"biasdseq"} = 1;

open(IN, $alt_list) || die "ERROR unable to open $alt_list for reading";
while(my $line = <IN>) { 
  ##     alert          short                            long       
  ##idx  code      S/F  description                      description
  ##---  --------  ---  -------------------------------  -----------
  #1     incsbgrp  S    Incorrect_Specified_Subgroup     score difference too large between best overall model and best specified subgroup model
  if($line !~ m/^\#/) { 
    chomp $line;
    my @el_A = split(/\s+/, $line);
    if(scalar(@el_A) < 5) { die "ERROR unable to parse line $line"; }
    ($code, $desc) = ($el_A[1], $el_A[3]);
    push(@code_A, $code);
    $desc_H{$code} = $desc;
  }
}
close(IN);

# make sure we read all non-fatal alerts
foreach $code (sort keys %nonfatal_H) { 
  if(! defined $desc_H{$code}) { die "ERROR did not read alt_list line for hard-coded non-fatal alert $code"; }
}

# get counts and num seqs for each code
my %seq_HH = ();
my %count_HH = ();
my %total_count_H = ();
my %total_seq_H = ();
my $count;
my $seq;
my %vapid_fail_H = ();
my %vapid_total_H = ();
my %vigor_fail_H = ();
my %vigor_total_H = ();
my $seqlist;

my %full_nseq_H = ();
my $full_nseq = 0;
foreach my $set (@sets_A) { 
  my $cur_nseq = undef;
  if($do_full) { 
    $cur_nseq = `grep -v ^\# $results_dir/va-$set/va-$set.vadr.sqc | wc -l`;
  }
  else { 
    $cur_nseq = `grep -v ^\# $results_dir/va-$set.r200/va-$set.r200.vadr.sqc | wc -l`;
  }
  chomp $cur_nseq;
  $full_nseq_H{$set} = $cur_nseq;
  $full_nseq += $cur_nseq;
}

foreach $code (@code_A) { 
  $vapid_fail_H{$code} = 0;
  $vapid_total_H{$code} = 0;
  $vigor_fail_H{$code} = 0;
  $vigor_total_H{$code} = 0;
  foreach my $set (@sets_A) { 
    if($do_full) { 
      $count = `grep -v ^\# $results_dir/va-$set/va-$set.vadr.alt | grep " $code " | wc -l`;
      $seq   = `grep -v ^\# $results_dir/va-$set/va-$set.vadr.alt | grep " $code " | awk '{ print \$2 }' | uniq | wc -l`;
    }
    else { 
      $count = `grep -v ^\# $results_dir/va-$set.r200/va-$set.r200.vadr.alt | grep " $code " | wc -l`;
      $seq   = `grep -v ^\# $results_dir/va-$set.r200/va-$set.r200.vadr.alt | grep " $code " | awk '{ print \$2 }' | uniq | wc -l`;
    }
    chomp $count;
    chomp $seq;
    #printf("$code $set $count $seq\n");
    $count_HH{$code}{$set} = $count;
    $seq_HH{$code}{$set} = $seq;
    $total_count_H{$code}  += $count;
    $total_seq_H{$code} += $seq;

    # if (! $do_full), compare with vapid and vigor
    # get a list of all seqs with that code
    if(! $do_full) { 
      my $seqlist = `grep -v ^\# $results_dir/va-$set.r200/va-$set.r200.vadr.alt | grep " $code " | awk '{ print \$2 }' | uniq`;
      my @seq_A = split(/\n/, $seqlist);
      foreach my $cur_seq (@seq_A) { 
        if(($set eq "nc") || ($set eq "np")) { 
          my $vigor_fail = `grep $cur_seq $set*combseqsum | grep vigor:FAIL | wc -l`;
          chomp $vigor_fail;
          if($vigor_fail eq "1") { $vigor_fail_H{$code}++; }
          $vigor_total_H{$code}++;
        }
        if(($set eq "nc") || ($set eq "dc")) { 
          my $vapid_fail = `grep $cur_seq $set*combseqsum | grep vapid:FAIL | wc -l`;
          chomp $vapid_fail;
          if($vapid_fail eq "1") { $vapid_fail_H{$code}++; }
          $vapid_total_H{$code}++;
        }
      }
    }
  }
}

# sort alert codes by total count
my @sorted_code_A = sort ({$total_count_H{$b} <=> $total_count_H{$a}} keys %total_count_H);
my @zero_ct_A = ();
my $nnonzero = 0;

# determine alerts with 0 instances
foreach $code (@sorted_code_A) { 
  if(! defined $nonfatal_H{$code}) { 
    if($total_count_H{$code} > 0) { 
      $nnonzero++;
    }
    else { 
      push(@zero_ct_A, $code); 
    }
  }
}
my $nzero = scalar(@zero_ct_A);

# output
print("\\begin{table}[h!]\n");
printf("\\caption{Counts of fatal VADR alerts reported for the test datasets. The $nnonzero fatal alert codes reported at least once for any test dataset are listed sorted by total number of reports. $nzero fatal alert types (");
for(my $c = 0; $c < ($nzero-2); $c++) { print(" $zero_ct_A[$c],"); }
print(" " . $zero_ct_A[$nzero-2] . " and " . $zero_ct_A[$nzero-1] . ". ");
print(") were not reported for any of the four test sets and are not shown.");
#printf("\$\^\*\$: these two alerts are non-fatal, all others are fatal (fatal alerts cause the corresponding sequence to fail). ");
if(! $do_full) { 
  print("The two rightmost columns indicate the numbers of sequences with the corresponding VADR alert that pass/fail VAPiD and VIGOR, respectively. Because VAPiD was tested on only NC (Norovirus-Complete) and DC (Dengue-Complete) and VIGOR was tested on only NC and NP (Norovirus-Partial), the numbers of sequences counted for VADR, VAPiD, and VIGOR differ on most rows of the table.\n");
}
printf("See Table~\\ref{tbl:alertlist} for more information on alerts.}\n");
if($do_full) { 
  print("\\begin{tabular}{|l|l||r|r|r|r||r|}\n");
  printf("\\hline\n");
  printf("         &                                     & NC      & NP      & DC      & DP      & total \\\\\n");
  printf(" alert   &                                     & %10s seqs &  %10s seqs &  %10s seqs &  %10s seqs & %10s seqs \\\\\n", $full_nseq_H{"nc"}, $full_nseq_H{"np"}, $full_nseq_H{"dc"}, $full_nseq_H{"dp"}, $full_nseq);
  printf(" code    & error message             & ct(seqs) & ct(seqs) & ct(seqs) & ct(seqs) & ct(seqs) \\\\ \\hline\n");
}
else { 
  print("\\begin{tabular}{|l|l||r|r|r|r||r||r||r|}\n");
  printf("\\hline\n");
  printf(" alert   &                                     & NC      & NP      & DC      & DP      & total    & VAPiD & VIGOR \\\\\n");
  printf(" code    & error message             & ct(seqs) & ct(seqs) & ct(seqs) & ct(seqs) & ct(seqs) & pass/fail & pass/fail \\\\ \\hline\n");
}
foreach $code (@sorted_code_A) { 
  if(! defined $nonfatal_H{$code}) { 

    my $desc2print = $desc_H{$code};
    $desc2print =~ s/\_/\\\_/g;
    my $code2print = $code;

    if($total_count_H{$code} > 0) { 
      printf("%13s & %-35s ", $code2print, $desc2print);
      foreach my $set (@sets_A) { 
        printf("& %12s ", ($count_HH{$code}{$set} == 0) ? "-" : sprintf("%d(%d)", $count_HH{$code}{$set}, $seq_HH{$code}{$set})); 
      }
      printf("& %12s ", sprintf("%d(%d)", $total_count_H{$code}, $total_seq_H{$code})); 

      if(! $do_full) { 
        # not doing full, output without fractions and with vigor and vapid totals
        if($vapid_total_H{$code} != 0) { 
          printf("& %8s ", sprintf("%d/%d", $vapid_total_H{$code} - $vapid_fail_H{$code}, $vapid_fail_H{$code}));
        }
        else { 
          printf("& %8s ", "-");
        }
        if($vigor_total_H{$code} != 0) { 
          printf("& %8s ", sprintf("%d/%d", $vigor_total_H{$code} - $vigor_fail_H{$code}, $vigor_fail_H{$code}));
        }
        else { 
          printf("& %8s ", "-");
        }
      }
      printf(" \\\\\n");
    }
  }
}
print("\\hline \n");
print("\\end{tabular}\n");
print("\\label{tbl:alertcounts}\n");
print("\\end{table}\n");
```
