## Supplemental material tarball for "VADR: validation and annotation of virus sequence submissions to GenBank": vadr-make-alertlist-table.pl

```
#!/usr/bin/env perl
use strict;

my $usage = "perl vadr-make-alertlist-table.pl <v-annotate.pl --alt_list output>";

if(scalar(@ARGV) != 1) { 
  die $usage;
}
my ($alt_list) = (@ARGV);

# hard-coded non-fatal alerts
my %nonfatal_H = ();
$nonfatal_H{"qstsbgrp"} = 1;
$nonfatal_H{"qstgroup"} = 1;
$nonfatal_H{"indfclas"} = 1;
$nonfatal_H{"lowscore"} = 1;
$nonfatal_H{"biasdseq"} = 1;

# hard-coded always fatal alerts
my %alwaysfatal_H = ();
$alwaysfatal_H{"noannotn"} = 1;
$alwaysfatal_H{"revcompl"} = 1;
$alwaysfatal_H{"unexdivg"} = 1;
$alwaysfatal_H{"noftrann"} = 1;

# hard-coded stage information
my %stage_H = ();
$stage_H{"noannotn"} = "classification";
$stage_H{"revcompl"} = "classification";
$stage_H{"unexdivg"} = "annotation";
$stage_H{"noftrann"} = "annotation";
$stage_H{"incsbgrp"} = "classification";
$stage_H{"incgroup"} = "classification";
$stage_H{"lowcovrg"} = "coverage";
$stage_H{"biasdseq"} = "coverage";
$stage_H{"dupregin"} = "coverage";
$stage_H{"discontn"} = "coverage";
$stage_H{"indfstrn"} = "coverage";
$stage_H{"lowsim5s"} = "coverage";
$stage_H{"lowsim3s"} = "coverage";
$stage_H{"lowsimis"} = "coverage";
$stage_H{"mutstart"} = "annotation";
$stage_H{"mutendcd"} = "annotation";
$stage_H{"mutendns"} = "annotation";
$stage_H{"mutendex"} = "annotation";
$stage_H{"unexleng"} = "annotation";
$stage_H{"cdsstopn"} = "annotation";
$stage_H{"cdsstopp"} = "protein validation";
$stage_H{"peptrans"} = "annotation";
$stage_H{"pepadjcy"} = "annotation";
$stage_H{"indfantp"} = "protein validation";
$stage_H{"indfantn"} = "annotation";
$stage_H{"indf5gap"} = "annotation";
$stage_H{"indf5loc"} = "annotation";
$stage_H{"indf5plg"} = "protein validation";
$stage_H{"indf5pst"} = "protein validation";
$stage_H{"indf3gap"} = "annotation";
$stage_H{"indf3loc"} = "annotation";
$stage_H{"indf3plg"} = "protein validation";
$stage_H{"indf3pst"} = "protein validation";
$stage_H{"indfstrp"} = "protein validation";
$stage_H{"insertnp"} = "protein validation";
$stage_H{"deletinp"} = "protein validation";
$stage_H{"lowsim5f"} = "annotation";
$stage_H{"lowsim3f"} = "annotation";
$stage_H{"lowsimif"} = "annotation";
$stage_H{"qstsbgrp"} = "classification";
$stage_H{"qstgroup"} = "classification";
$stage_H{"indfclas"} = "classification";
$stage_H{"lowscore"} = "classification";

my $code;
my $desc;
my $long;
my $per;
my @code_A = ();
my %desc_H = ();
my %long_H = ();
my %per_H  = ();

open(IN, $alt_list) || die "ERROR unable to open $alt_list for reading";
while(my $line = <IN>) { 
  ##     alert          short                            long       
  ##idx  code      S/F  description                      description
  ##---  --------  ---  -------------------------------  -----------
  #1     incsbgrp  S    Incorrect_Specified_Subgroup     score difference too large between best overall model and best specified subgroup model
  if($line !~ m/^\#/) { 
    chomp $line;
    my @el_A = split(/\s+/, $line);
    if(scalar(@el_A) < 5) { die "ERROR unable to parse line $line"; }
    ($code, $per, $desc) = ($el_A[1], $el_A[2], $el_A[3]);
    push(@code_A, $code);
    $desc_H{$code} = $desc;
    $per_H{$code} = $per;
    if($line =~ /^\d+\s+\S+\s+\S+\s+\S+\s+(.+)$/) { 
      $long_H{$code} = $1;
    }
    else { 
      die "ERROR couldn't parse long desc from line $line";
    }
  }
}
close(IN);

# make sure we read all alert codes
foreach $code (sort keys %stage_H) { 
  if(! defined $desc_H{$code}) { die "ERROR did not read alt_list line for hard-coded alert $code"; }
}
foreach $code (@code_A) { 
  if(! defined $stage_H{$code}) { die "ERROR read alert we don't have stage info for: $code"; }
}
foreach $code (sort keys %nonfatal_H) { 
  if(! defined $desc_H{$code}) { die "ERROR did not read alt_list line for hard-coded non-fatal alert $code"; }
}
foreach $code (sort keys %alwaysfatal_H) { 
  if(! defined $desc_H{$code}) { die "ERROR did not read alt_list line for hard-coded always-fatal alert $code"; }
}

my $stage;
# output
print("\\begin{table}[h!]\n");
printf("\\caption{Attributes of the 43 types of VADR alerts organized by the stage in which they are detected. The S/F column indicates whether the alert applies to an entire sequence (S) or to one feature (F) in a sequence. The five non-fatal alerts (four detected in the classification stage, and one in the coverage stage) do not cause a sequence to fail and are not reported in the output feature table. Codes marked with \$^{*}\$ are always fatal; all other codes can be set to fatal or non-fatal with command-line options to v-annotate.pl.}\n");
print("\\begin{tabular}{|l|l|l|l|}\n");
printf("\\hline\n");
printf("\\textbf{code} & \\textbf{S/F} & \\textbf{error message} & \\textbf{description}  \\\\\n");

my $nprinted = 0;
my $long_maxlen = 65;
foreach $stage ("classification", "coverage", "annotation", "protein validation") { 
  $nprinted = 0;
  # fatal alerts
  foreach $code (@code_A) { 
    my $code2print = $code;
    if(defined $alwaysfatal_H{$code}) { 
      $code2print .= "\$^{*}\$";
    }
    my $desc2print = $desc_H{$code};
    my $long2print = $long_H{$code};
    $desc2print =~ s/\_/\\\_/g;
    $long2print =~ s/\_/\\\_/g;

    # split up long descriptions over two lines
    my $long2print1 = $long2print;
    my $line2 = undef;
    if(length($long2print) > $long_maxlen) { 
      # find first space or dash after $long_maxlen to split on
      my $cur_maxlen = $long_maxlen;
      while(($cur_maxlen < length($long2print)) && (substr($long2print, $cur_maxlen, 1) !~ m/[ \-]/)) { 
        $cur_maxlen++;
      }
      if($cur_maxlen < length($long2print)) { 
        $long2print1 = substr($long2print, 0, $cur_maxlen);
        my $long2print2 = substr($long2print, $cur_maxlen);
        $line2 = sprintf("%-14s & %1s & %-32s & %s \\\\\n", "", "", "", $long2print2);
      }
    }
    
    if(($stage_H{$code} eq $stage) && (! defined $nonfatal_H{$code})) { 
      if($nprinted == 0){ 
        printf("\\hline \n");
        printf("\\multicolumn{4}{|l|}{\\textbf{Fatal alerts detected in the $stage stage}} \\\\\n");
        printf("\\hline \n");
      }
      printf("%-14s & %1s & %-32s & %s \\\\\n", $code2print, $per_H{$code}, $desc2print, $long2print1);
      if(defined $line2) { print $line2; }
      $nprinted++;
    }
  }
  # non-fatal alerts
  $nprinted = 0;
  foreach $code (@code_A) { 
    my $code2print = $code;
    if(defined $alwaysfatal_H{$code}) { 
      $code2print .= "\$^{*}\$";
    }
    my $desc2print = $desc_H{$code};
    my $long2print = $long_H{$code};
    $desc2print =~ s/\_/\\\_/g;
    $long2print =~ s/\_/\\\_/g;

    # split up long descriptions over two lines
    my $long2print1 = $long2print;
    my $line2 = undef;
    if(length($long2print) > $long_maxlen) { 
      # find first space or dash after $long_maxlen to split on
      my $cur_maxlen = $long_maxlen;
      while(($cur_maxlen < length($long2print)) && (substr($long2print, $cur_maxlen, 1) !~ m/[ \-]/)) { 
        $cur_maxlen++;
      }
      if($cur_maxlen < length($long2print)) { 
        $long2print1 = substr($long2print, 0, $cur_maxlen);
        my $long2print2 = substr($long2print, $cur_maxlen);
        $line2 = sprintf("%-14s & %1s & %-32s & %s \\\\\n", "", "", "", $long2print2);
      }
    }

    if(($stage_H{$code} eq $stage) && (defined $nonfatal_H{$code})) { 
      if($nprinted == 0){ 
        printf("\\hline \n");
        printf("\\multicolumn{4}{|l|}{\\textbf{Non-fatal alerts detected in the $stage stage}} \\\\\n");
        printf("\\hline \n");
      }
      printf("%-14s & %1s & %-32s & %s \\\\\n", $code2print, $per_H{$code}, $desc2print, $long2print1);
      if(defined $line2) { print $line2; }
      $nprinted++;
    }
  }
#  printf(" & & & \\\\\n");
}
print("\\\hline \n");
print("\\end{tabular}\n");
print("\\label{tbl:alertlist}\n");
print("\\end{table}\n");
```
