## Supplemental material tarball for "VADR: validation and annotation of virus sequence submissions to GenBank": vadr-make-full-results-table.pl

```
if(! exists($ENV{"VADRPAPER_VADRDIR"})) { 
  die "ERROR, the environment variable VADRPAPER_VADRDIR is not set";
}
my $dir = $ENV{"VADRPAPER_VADRDIR"} . "/vadr-full-out";
if(! -d $dir) { die "ERROR results dir $dir does not exist"; }

my @key_A = ("nc", "dc", "np", "dp");

foreach $key (@key_A) { 
  my $npass = `grep -v ^\# $dir/va-$key/va-$key*.sqa | grep " PASS " | wc -l`;
  my $nfail = `grep -v ^\# $dir/va-$key/va-$key*.sqa | grep " FAIL " | wc -l`;
  chomp $npass;
  chomp $nfail;
  $npass_H{$key} = $npass;
  $nfail_H{$key} = $nfail;
  #printf("$key $npass_H{$key} $nfail_H{$key}\n");
}

print("\\begin{table}[t]\n");
print("\\caption{Summary of pass/fail outcomes for VADR on the full datasets}\n");
print("\\begin{tabular}{|l|r|}\n");
############################
printf("\\hline\n");
printf("         & \\multicolumn{1}{c|}{VADR}       \\\\\n");
printf(" dataset & \\multicolumn{1}{c|}{\\#pass/\\#fail [pass fraction]}  \\\\ \\hline\n");
foreach $key (@key_A) { 
  my $key2print = $key;
  $key2print =~ tr/a-z/A-Z/;
  if($key2print eq "NA") { $key2print = "NP"; }
  if($key2print eq "DA") { $key2print = "DP"; }
  #printf("%9s ", "\\emph\{" . $key12print . "\}");
  printf("%9s ", $key2print);
  printf("& %7s [%.3f]", $npass_H{$key} . "/" . $nfail_H{$key}, ($npass_H{$key}/ ($npass_H{$key} + $nfail_H{$key})));
  print("\\\\ \n");
}
printf("\\hline \n");
############################
print("\\end{tabular}\n");
print("\\label{tbl:fullpassfail}\n");
print("\\end{table}\n");
```
