## Supplemental material tarball for "VADR: validation and annotation of virus sequence submissions to GenBank": vadr-make-results-tables.pl

```
while($line = <>) { 
  chomp $line;
  if($line =~ m/\w/) { 
    @el_A = split(/\s+/, $line);
    if(scalar(@el_A) != 4) { die "ERROR unable to parse line $line"; }
    $count_HHH{$el_A[0]}{$el_A[1]}{$el_A[2]} = $el_A[3];
  }
}


print("\\begin{table}[t]\n");
print("\\caption{Summary of pass/fail outcomes for VADR, VAPiD and VIGOR on the 200 sequence test datasets}\n");
print("\\begin{tabular}{|l|r|r|r|}\n");
############################
printf("\\hline\n");
printf("         & VADR       & VAPiD     & VIGOR      \\\\\n");
printf(" dataset & pass/fail  & pass/fail & pass/fail  \\\\ \\hline\n");
foreach $key1 ("nc", "dc", "np", "dp") { 
  my $key12print = $key1;
  $key12print =~ tr/a-z/A-Z/;
  #printf("%9s ", "\\emph\{" . $key12print . "\}");
  printf("%9s ", $key12print);
  foreach $key2 ("vadr", "vapid", "vigor") { 
    if($count_HHH{$key1}{$key2}{"pass"} eq "-") { 
      printf("& %7s ", "-");
    }
    else { 
      printf("& %7s ", $count_HHH{$key1}{$key2}{"pass"} . "/" . $count_HHH{$key1}{$key2}{"fail"});
    }
  }
  print("\\\\ \n");
}
printf("\\hline \n");
############################
print("\\end{tabular}\n");
print("\\label{tbl:passfail}\n");
print("\\end{table}\n");

#======================================================================
printf("\n\n\n");

print("\\begin{table}[t]\n");
print("\\caption{Comparison of pass/fail outcomes for VADR and VAPiD on the 200 sequence Norovirus-Complete (NC) and Dengue-Complete (DC) test datasets.}\n");
print("\\begin{tabular}{|l|r|r|r|r|}\n");
############################
printf("\\hline\n");
printf("         & Both       & Both       & VADR-pass  & VADR-fail  \\\\\n");
printf(" dataset & pass       & fail       & VAPiD-fail & VAPiD-pass \\\\ \\hline\n");
foreach $key1 ("nc", "dc") { 
  my $key12print = $key1;
  $key12print =~ tr/a-z/A-Z/;
#  printf("%9s ", "\\emph\{" . $key12print . "\}");
  printf("%9s ", $key12print);
  $key2 = "vadr-vapid";
  foreach $key3 ("pass-pass", "fail-fail", "pass-fail", "fail-pass") { 
    printf("& %6s ", $count_HHH{$key1}{$key2}{$key3}); 
  }
  print("\\\\ \n");
}
printf("\\hline \n");
############################
print("\\end{tabular}\n");
print("\\label{tbl:compare-vadr-vapid}\n");
print("\\end{table}\n");

############################
printf("\n\n\n");
############################

print("\\begin{table}[t]\n");
print("\\caption{Comparison of pass/fail outcomes for VADR and VIGOR on the 200 sequence Norovirus-Complete (NC) and Norovirus-Partial (NP) test datasets.}\n");
print("\\begin{tabular}{|l|r|r|r|r|}\n");
############################
printf("\\hline\n");
printf("         & Both       & Both       & VADR-pass  & VADR-fail  \\\\\n");
printf(" dataset & pass       & fail       & VIGOR-fail & VIGOR-pass \\\\ \\hline\n");
foreach $key1 ("nc", "np") { 
  my $key12print = $key1;
  $key12print =~ tr/a-z/A-Z/;
#  printf("%9s ", "\\emph\{" . $key12print . "\}");
  printf("%9s ", $key12print);
  $key2 = "vadr-vigor";
  foreach $key3 ("pass-pass", "fail-fail", "pass-fail", "fail-pass") { 
    printf("& %6s ", $count_HHH{$key1}{$key2}{$key3}); 
  }
  print("\\\\ \n");
}
printf("\\hline \n");
############################
print("\\end{tabular}\n");
print("\\label{tbl:compare-vadr-vigor}\n");
print("\\end{table}\n");
```
