## Supplemental material tarball for "VADR: validation and annotation of virus sequence submissions to GenBank": vadr-sqa-and-ftr-to-seqsum-and-ftrsum.pl

```
#!/usr/bin/env perl
use strict;

my $usage = "perl vadr-sqa-and-ftr-to-seqsum-and-ftrsum.pl <VADR .sqa.tbl output file> <VADR .ftr.tbl output file> <map file with bestmdl:cds-product:protien-acc info> <output root, e.g. \"vadr.np.\">";

if(scalar(@ARGV) != 4) { die $usage; }

my ($sqa_file, $ftr_file, $map_file, $out_root) = (@ARGV);

my $line; # a line of the .at file
my %refprotein_H = ();
my %refseq_exists_H = ();
# parse the map file
open(MAP, $map_file) || die "ERROR unable to open $map_file for reading";
while($line = <MAP>) { 
#NC_001959:nonstructural polyprotein:NP_056820.1
  chomp $line;
  if($line =~ m/^([^\:]+)\:([^\:]+)\:([^\:]+)$/) { 
    my ($refseq, $product, $refprotein) = ($1, $2, $3);
    $product =~ s/\s/\_/g;
    $refseq_exists_H{$refseq} = 1;
    $refprotein_H{($refseq . ":" . $product)} = $refprotein;
  }
}

my $out_seqsum_file = $out_root . ".seqsum";
my $out_ftrsum_file = $out_root . ".ftrsum";
open(SEQSUM, ">", $out_seqsum_file) || die "ERROR unable to open $out_seqsum_file for writing";
open(FTRSUM, ">", $out_ftrsum_file) || die "ERROR unable to open $out_ftrsum_file for writing";

my ($seqidx, $seqname, $seqlen, $pf, $ant, $bestmdl, $grp, $subgrp, $nfa, $nfn, $nf5, $nf3, $nfalt, $seqalerts);
my $line; # a line of the .at file
my %pf_H = (); # key is sequence name, value is PASS or FAIL
my %cls_H = (); # key is sequence name, value is $grp:$subgrp
my %pfreason_H = (); # key is sequence name, value is reason for FAILure or "NULL" if it passes
my %seqname_exists_H = (); # key is sequence name, value is '1'
my @seqname_A = (); # array of sequence names in order they were read in

open(SQA, $sqa_file) || die "ERROR unable to open $sqa_file for reading";
while($line = <SQA>) { 
  if($line !~ m/^\#/) { 
    chomp $line;
    my @el_A = split(/\s+/, $line);
    if(scalar(@el_A) == 14) { 
      ##seq  seq           seq             best               sub                             seq   
      ##idx  name          len  p/f   ant  model      grp     grp  nfa  nfn  nf5  nf3  nfalt  alerts
      ###---  ----------  -----  ----  ---  ---------  ------  ---  ---  ---  ---  ---  -----  ------
      # 1     HQ541796.1  10539  PASS  yes  NC_001475  Dengue  3     24    2    1    2      0  -     
      ($seqidx, $seqname, $seqlen, $pf, $ant, $bestmdl, $grp, $subgrp, $nfa, $nfn, $nf5, $nf3, $nfalt, $seqalerts) = @el_A;
      
      push(@seqname_A, $seqname);
      $pf_H{$seqname}  = $pf;
      $cls_H{$seqname} = $grp . ":" . $subgrp;

      if($seqalerts ne "-") { 
        # have to look at the seqalerts to skip those that are not fatal
        my @alert_A = split(",", $seqalerts);
        my $alerts_to_add = "";
        foreach my $alert_to_add (@alert_A) { 
          if($alert_to_add !~ m/\(qstsbgrp\)/ && 
             $alert_to_add !~ m/\(qstgroup\)/ && 
             $alert_to_add !~ m/\(indfclas\)/ && 
             $alert_to_add !~ m/\(lowscore\)/) { 
            # skip non-fatal alerts
            if($alerts_to_add ne "") { $alerts_to_add .= ","; }
            $alerts_to_add .= $alert_to_add;
          }
        }
        $pfreason_H{$seqname} = $alerts_to_add;
      }
      else { 
        $pfreason_H{$seqname} = "";
      }
    }
    else { 
      die "ERROR unable to parse $line";
    }
  }
}
close(SQA);

open(FTR, $ftr_file) || die "ERROR unable to open $ftr_file for reading";
my ($idx, $ftrtype, $ftrname, $ftrlen, $ftridx, $str, $n_from, $n_to, $n_instp, $trc, $p_from, $p_to, $p_instp, $p_sc, $nsa, $nsn, $seq_coords, $mdl_coords, $ftr_alerts);
while($line = <FTR>) { 
  if($line !~ m/^\#/) { 
    chomp $line;
    my @el_A = split(/\s+/, $line);
    if(scalar(@el_A) == 23) { 
      ($idx, $seqname, $seqlen, $pf, $bestmdl, $ftrtype, $ftrname, $ftrlen, $ftridx, $str, $n_from, $n_to, $n_instp, $trc, $p_from, $p_to, $p_instp, $p_sc, $nsa, $nsn, $seq_coords, $mdl_coords, $ftr_alerts) = (@el_A);
      
      if(! defined $pf_H{$seqname}) { die "ERROR read sequence $seqname in ftr file that was not in sqa file"; }
      if($pf_H{$seqname} ne $pf) { die "ERROR pass fail disagreement for $seqname"; }

      if($ftr_alerts ne "-") { 
        if($pfreason_H{$seqname} ne "") { $pfreason_H{$seqname} .= ","; }
        $pfreason_H{$seqname} .= $ftr_alerts;
      }

      # determine reference protein
      my $refprotein = "IRRELEVANT";
      if(($ftrtype eq "CDS") && (defined $refseq_exists_H{$bestmdl})) { 
        my $tmpkey = $bestmdl . ":" . $ftrname;
        if(! defined $refprotein_H{$tmpkey}) { 
          die "ERROR no reference protein for $tmpkey";
        }
        $refprotein = $refprotein_H{$tmpkey};
      }
      printf FTRSUM ("seq:$seqname product:$ftrname ref:$refprotein coords:$seq_coords pf:$pf_H{$seqname}\n");
    }
  }
}
close(FTR);

foreach my $seqname (@seqname_A) { 
  if($pfreason_H{$seqname} eq "") { 
    $pfreason_H{$seqname} = "NULL";
  }
  printf SEQSUM ("seq:%-10s pf:%s pfreason:%s cls:%s\n", $seqname, $pf_H{$seqname}, $pfreason_H{$seqname}, $cls_H{$seqname});
}

close(SEQSUM);
close(FTRSUM);

exit 0;
```
