## Supplemental material tarball for "VADR: validation and annotation of virus sequence submissions to GenBank": vadr-vigor-compare-ftrsums.sh

```
#!/bin/bash
for p in all pass; do 
    for a in np nc; do 
        echo $a
        perl $VADRPAPER_SCRIPTSDIR/compare-two-ftrsums.pl vadr.$a.ftrsum vigor.$a.ftrsum $p > vadr.vigor.$a.$p.compftrsum
        echo -n "$p count of PRODUCT-MATCH "
        cat vadr.vigor.$a.$p.compftrsum | grep PRODUCT-MATCH | wc -l
        echo -n "$p count of PRODUCT-MATCH COORDS-MATCH "
        cat vadr.vigor.$a.$p.compftrsum | grep PRODUCT-MATCH | grep COORDS-MATCH | wc -l
        echo -n "$p count of PRODUCT-MATCH COORDS-MISMATCH "
        cat vadr.vigor.$a.$p.compftrsum | grep PRODUCT-MATCH | grep COORDS-MISMATCH | wc -l
        echo ""
    done
    echo "$p np and nc combined"
    echo -n "$p count of PRODUCT-MATCH "
    cat vadr.vigor.n*.$p.compftrsum | grep PRODUCT-MATCH | wc -l
    echo -n "$p count of PRODUCT-MATCH COORDS-MATCH "
    cat vadr.vigor.n*.$p.compftrsum | grep PRODUCT-MATCH | grep COORDS-MATCH | wc -l
    echo -n "$p count of PRODUCT-MATCH COORDS-MISMATCH "
    cat vadr.vigor.n*.$p.compftrsum | grep PRODUCT-MATCH | grep COORDS-MISMATCH | wc -l
    echo ""
done
```
