## Supplemental material tarball for "VADR: validation and annotation of virus sequence submissions to GenBank": vapid-out-and-errorsummary-to-seqsum.pl

```
#!/usr/bin/env perl
use strict;

my $usage = "perl vapid-out-and-errorsummary-to-seqsum.pl\n\t<vapid .out output file>\n\t<dir with possible */errorsummary.val file>\n\t<namelist file with name of sequence .out file corresponds to>";

if(scalar(@ARGV) != 3) { die $usage; }

my ($infile, $errdir, $namefile) = (@ARGV);
my $short_infile = $infile;
$short_infile =~ s/^.+\///;

open(NAME, $namefile) || die "ERROR unable to open $namefile for reading";
my $namefile_seqname = <NAME>;
chomp $namefile_seqname;
close(NAME);


# determine path to errorsummary.val file, if it exists
my $errsubdir = $errdir . "/" . $namefile_seqname;
if(! -d $errsubdir) { die "ERROR error subdir $errsubdir does not exist"; }
my $errfile = $errsubdir . "/errorsummary.val";

open(IN, $infile) || die "ERROR unable to open $infile for reading";

#~~~~~~~~~~~~~~~~~~~~~~~~
#Automatic strain naming failed but submission will proceed without metadata appended to the fasta header.
#Searching local blast database at all_virus.fasta
#GQ868627.1 was the selected reference
#Dengue virus 3 was the parsed name of the virus
#Aligning reference and query...
#Done alignment
#polyprotein ['74', '10246']
#Done with: GQ868627.1
#
#
#Done, did  1 viruses in 3.4 seconds
#~~~~~~~~~~~~~~~~~~~~~~~~

my $seqname = undef;
my $warn_seqname = undef;
my $refname = undef;
my $hasstop = undef;
my $nstop = undef;
my $revcomp = 0;
my $seconds = undef;

while(my $line = <IN>) { 
  chomp $line;
  if($line =~ /^Searching local blast database at (\S+)/) { 
    if($1 ne "all_virus.fasta") { 
      die "ERROR unexpected local blast db:\n$line\n";
    }
  }
  elsif($line =~ /^(\S+) was the selected reference/) { 
    if(defined $refname) { 
      if(! $revcomp) { 
        die "ERROR more than one reference line (and not reverse complemented)"; 
      }
    }
    $refname = $1;
  }
  elsif($line =~ /^Done with\:\s+(\S+)/) { 
    if(defined $seqname) { die "ERROR more than one 'Done with' line (and not reverse complemented)"; }
    $seqname = $1;
    if($seqname ne $namefile_seqname) { 
      die "ERROR sequence name read doesn't match name in namelist file $seqname ne $namefile_seqname"; 
    }
  }
  elsif($line =~ /^Input sequence needed to be reverse complemented to align properly\.$/) { 
    $revcomp = 1; 
  }
  elsif($line =~ /^WARNING\:\ (\S+) contains (\d+) stop codon\(s\)\!$/) { 
    $warn_seqname = $1;
    $hasstop = 1;
    $nstop = $2;
    if(! defined $seqname) { die "ERROR read WARNING line before 'Done with' line"; }
    if($warn_seqname ne $seqname) { die "ERROR sequence name on WARNING line doesn't match seq name from 'Done with' line"; }
  }
  elsif($line =~ /^WARNING/) { 
    die "ERROR unexpected WARNING line: $line\n";
  }
  elsif($line =~ /^Done, did  1 viruses in (\S+) seconds$/) {
    $seconds = $1;
  }
  # and make sure you don't see any unexpected lines:
  elsif($line =~ /^Automatic strain naming failed but submission will proceed without metadata appended to the fasta header\.$/) { 
    ; # do nothing
  }
  elsif($line =~ /^.+was the parsed name of the virus$/) { 
    ; # do nothing
  }
  elsif($line =~ /^Aligning reference and query\.\.\.$/) { 
    ; # do nothing
  }
  elsif($line =~ /^Done alignment$/) { 
    ; # do nothing
  }
  elsif($line =~ /^.+\[\'?(\-?\d+)\'?\,\s+\'?(\-?\d+)\'?\]$/) { 
    my ($start, $stop) = ($1, $2);
  }
  elsif($line eq "") { 
    ; # do nothing
  }
  elsif($line =~ /^Modifying ORF length for/) { 
    ; # do nothing
  }
  else { 
    die "ERROR unexpected line:\n$line\n";
  }
}
close(IN);

# does this sequence find a reference? Or are there zero suitable blast hits?
my $no_ref = 0; # set to 1 if there is no reference (no blast hit)
if(! defined $seqname) { 
  $no_ref = 1;
  $seqname = $namefile_seqname;
  $refname = "UNKNOWN";
}  

# 
if(! defined $hasstop) { 
  $hasstop = 0; 
  $nstop = 0; 
}

# we should have a time unless there's no reference
if(! defined $seconds) { 
  if($no_ref == 0) { 
    die "ERROR did not read timing line and reference is defined"; 
  }
  $seconds = "0.0";
}

my $ref_is_seq = 0;
if(($no_ref == 0) && ($seqname eq $refname)) { 
  $ref_is_seq = 1;
}

my $pfreason = "";
if($hasstop) { 
  if($pfreason ne "") { $pfreason .= ","; }
  $pfreason .= "nstop=$nstop"; 
}
if($no_ref) { 
  if($pfreason ne "") { $pfreason .= ","; }
  $pfreason .= "noref"; 
}
if($revcomp) { 
  if($pfreason ne "") { $pfreason .= ","; }
  $pfreason .= "revcomp"; 
}

# now process the errorsummary.val file if it exists
if(-e $errfile) { 
  if(-s $errfile) { 
    # parse it
    open(ERR, $errfile) || die "ERROR unable to open $errfile for reading"; 
    while(my $errline = <ERR>) { 
      chomp $errline;
      my $errval;
      if($errline =~ /^\s+\d+\s+ERROR\:\s+(\S+)$/) { 
        #1 ERROR:   SEQ_FEAT.NoStop
        $errval = "ERROR:" . $1;
      }
      elsif($errline =~ /^\s+\d+\s+WARNING\:\s+(\S+)$/) { 
        #2 WARNING: SEQ_INST.TerminalNs
        $errval = "WARNING:" . $1;
      }
      else { 
        die "ERROR unable to parse error line from $errfile:\n$errline\n"; 
      }
      if($pfreason eq "") { 
        $pfreason = $errval;
      }
      else { 
        $pfreason .= "," . $errval;
      }
    }
    close(ERR);
  }
  else { 
    ; # errfile exists and is empty, do nothing
  }
}
else { 
  # errfile does not exist
  # should only happen if $no_ref == 1
  if(! $no_ref) { die "ERROR errfile $errfile does not exist and no_ref is 0"; }
}

my $pf = undef;
if($pfreason eq "") { 
  $pfreason = "NULL";
  $pf = "PASS";
}
else { 
  $pf = "FAIL";
}

# output line
printf("seq:%-10s pf:$pf pfreason:$pfreason outfile:%-20s ref:%-10s ref_is_seq:%d no_ref:%d revcomp:%d nstop:%d seconds:%s\n", $seqname, $short_infile, $refname, $ref_is_seq, $no_ref, $revcomp, $nstop, $seconds);
```
