## Supplemental material tarball for "VADR: validation and annotation of virus sequence submissions to GenBank": vigor-at-to-seqsum-and-ftrsum.pl

```
#!/usr/bin/env perl
use strict;

my $usage = "perl vigor-at-to-seqsum-and-ftrsum.pl <VIGOR .at output file> <output root, e.g. \"vigor.np.\"";

if(scalar(@ARGV) != 2) { die $usage; }

my ($at_file, $out_root) = (@ARGV);

my $out_seqsum_file = $out_root . ".seqsum";
my $out_ftrsum_file = $out_root . ".ftrsum";
open(SEQSUM, ">", $out_seqsum_file) || die "ERROR unable to open $out_seqsum_file for writing";
open(FTRSUM, ">", $out_ftrsum_file) || die "ERROR unable to open $out_ftrsum_file for writing";

open(AT, $at_file) || die "ERROR unable to open $at_file for reading";
my ($seqname, $db, $coords, $refname, $gene, $product, $errcode, $errmsg); # tab-delimited tokens in the .at file
my $line; # a line of the .at file
my %product_H = (); # product value with possible 'truncated' and 'putative ' and 'fragment' and 'terminal' parts removed
my %pf_H = (); # key is sequence name, value is PASS or FAIL
my %db_H = (); # key is sequence name, value is db used for that sequence
my %line_beg_HA = (); # key is sequence name, value is array of first 6 tokens of all .at lines for that sequence
my %pfreason_H = (); # key is sequence name, value is reason for FAILure or "NULL" if it passes
my %line_beg_exists_H = (); # key is first 6 tokens of an .at file tab-delimited, value is '1'
my %seqname_exists_H = (); # key is sequence name, value is '1'
my @seqname_A = (); # array of sequence names in order they were read in

my %errcode_map_H = ();
$errcode_map_H{"FS"} = "FRAMESHIFT";
$errcode_map_H{"GS"} = "GAPERROR";
$errcode_map_H{"ES"} = "EMBEDDEDSTOP";
$errcode_map_H{"IS"} = "INVALIDSPLICING";
$errcode_map_H{"C5"} = "STARTCODON";
$errcode_map_H{"C3"} = "STOPCODON";

while($line = <AT>) { 
  chomp $line;
  my @el_A = split(/\t/, $line);
  my $line_beg = $el_A[0]."\t".$el_A[1]."\t".$el_A[2]."\t".$el_A[3]."\t".$el_A[4]."\t".$el_A[5];
  if(scalar(@el_A) == 8) { # should include an 'error': one of 'T5', 'T3', 'FS', 'GS', 'ES', 'IS'
    #AM905365.1	norv_misc_db	<1..>283	BAC98439.1	POL	putative nonstructural polyprotein, fragment	T5	4317
    #AM905365.1	norv_misc_db	<1..>283	BAC98439.1	POL	putative nonstructural polyprotein, fragment	T3	501
    ($seqname, $db, $coords, $refname, $gene, $product, $errcode, $errmsg) = (@el_A); 
    if(($errcode ne "T5") && 
       ($errcode ne "T3") && 
       ($errcode ne "FS") &&
       ($errcode ne "GS") && 
       ($errcode ne "ES") && 
       ($errcode ne "IS")) { 
      die "ERROR unexpected error code for 8 token line:\n$line\n";
    }
  }
  elsif(scalar(@el_A) == 7) { # 7th token (errcode) should be one of 'OK', 'C5' or 'C3'
    #KX353996.1	norv2_db	1..1623	AAZ31396.2	VP1	capsid protein VP1	OK	
    ($seqname, $db, $coords, $refname, $gene, $product, $errcode) = (@el_A); 
    if(($errcode ne "OK") && 
       ($errcode ne "C5") && 
       ($errcode ne "C3")) { 
      die "ERROR unexpected error code for 7 token line:\n$line\n";
    }
    $errmsg = ""; # no error message in 7 token lines
  }
  else { 
    die "ERROR read non-8 and non-7 token line: $line\n";
  }

  # product may have 'putative', and 'fragment/terminal', remove it so we can compare using equality with VADR output
  $product =~ s/^truncated //;
  $product =~ s/^putative //;
  $product =~ s/, fragment$//;
  $product =~ s/, C\-terminal$//;
  $product =~ s/, N\-terminal$//;

  $product_H{$line_beg} = $product;

  if(! defined $seqname_exists_H{$seqname}) { 
    # first line for this sequence
    push(@seqname_A, $seqname);
    $pf_H{$seqname} = "PASS";
    $seqname_exists_H{$seqname} = 1;
    @{$line_beg_HA{$seqname}} = ();
    $db_H{$seqname} = $db;
  }
  if(! defined $line_beg_exists_H{$line_beg}) { 
    push(@{$line_beg_HA{$seqname}}, $line_beg);
    $line_beg_exists_H{$line_beg} = 1;
  }
  if(($errcode ne "OK") && ($errcode ne "T5") && ($errcode ne "T3")) { 
    if(! defined $errcode_map_H{$errcode}) { die "ERROR no map for errcode $errcode"; }
    # this is a bad error that causes the sequence to FAIL
    $pf_H{$seqname} = "FAIL";
    if($pfreason_H{$seqname} ne "") { $pfreason_H{$seqname} .= ","; }
    $pfreason_H{$seqname} .= $errcode_map_H{$errcode} . "[" . $errmsg . "]";
  }
}
close(AT);

foreach my $seqname (@seqname_A) { 
  if(! defined $pfreason_H{$seqname}) { 
    $pfreason_H{$seqname} = "NULL";
  }
  printf SEQSUM ("seq:%-10s pf:%s pfreason:%s db:%s\n", $seqname, $pf_H{$seqname}, $pfreason_H{$seqname}, $db_H{$seqname});
  foreach my $line_beg (@{$line_beg_HA{$seqname}}) { 
    # parse the $line_beg into its tab-delimited tokens
    my @el_A = split(/\t/, $line_beg);
    if(scalar(@el_A) != 6) { die "ERROR line_beg did not have 6 tokens:\n$line_beg" };
    my ($seqname, $db, $coords, $refname, $gene, $product) = (@el_A);

    my $str = undef;
    my @coords_tok_A = split(",", $coords);
    my $new_coords = "";
    foreach my $coords_tok (@coords_tok_A) { 
      if($new_coords ne "") { $new_coords .= ","; }
      if($coords_tok =~ /^\<?(\d+)\.\.\>?(\d+)$/) { 
        my ($start, $stop) = ($1, $2);
        if   ($start < $stop)  { $new_coords .= $coords_tok . ":+"; }
        elsif($stop  < $start) { $new_coords .= $coords_tok . ":-"; }
        else { die "ERROR unable to parse strand from $coords"; }
      }
      else { 
        die "ERROR unable to parse coords_tok $coords_tok from line_beg: $line_beg";
      }
    }
    my $product2print = $product_H{$line_beg};
    $product2print =~ s/\s/\_/g;
    #printf FTRSUM ("seq:$seqname product:$product_H{$line_beg} gene:$gene coords:$new_coords pf:$pf_H{$seqname}\n");
    printf FTRSUM ("seq:$seqname product:$product2print coords:$new_coords ref:$refname pf:$pf_H{$seqname}\n");
  }
}
close(SEQSUM);
close(FTRSUM);

exit 0;
```
